# *In silico* pharmacological analysis of *Tinospora cordifolia* compounds targeting African swine fever virus B175L

**DOI:** 10.1101/2025.08.03.668349

**Authors:** Kasuni Karunarathne, Saumya Poorni, Nethra Jayampathi, Hasitha Dhananjaya, Anju Hasintha, W.M.M.P. Hulugalla, N.M.T. Anupama, Nadeeka Nethmini, Naveed Iqbal, Barana Jayawardana, Lakmal Ranathunga

**Affiliations:** Department of Animal Science, Faculty of Agriculture, University of Peradeniya, Peradeniya, Sri Lanka; Faculty of Veterinary Medicine and Animal Science, University of Peradeniya, Peradeniya, Sri Lanka; School of Dentistry and Medical Science, Charles Sturt University, NSW, Australia; School of Interdisciplinary Engineering and Sciences (SINES), National University of Sciences and Technology (NUST), Islamabad, Pakistan

## Abstract

African swine fever virus (ASFV) is a highly lethal DNA virus that suppresses the host’s immune response by establishing infection. B175L, one of its key immune evasion proteins, directly inhibits STING-mediated type I interferon (IFN-I) signaling, preventing antiviral defense activation. Thus, targeting B175L could be a promising strategy for antiviral drug development as effective ASFV inhibitors remain unidentified. In this study, we investigated the potential of the *Tinospora cordifolia* plant’s bioactive compounds to disrupt B175L’s function, restoring immune signaling. Gas chromatography-mass spectrometry (GC-MS) analysis of *T. cordifolia* stem’s methanol extract identified 124 compounds, of which 52 met SwissADME and DataWarrior criteria for drug-likeness and safety. We generated a highly accurate 3D B175L model with strong confidence scores for the structure accuracy, utilizing AlphaFold3. Virtual screening was performed using PyRx 0.8, and the top 4 ligands with binding affinities exceeding -6 kcal/mol advanced to CB-Dock2 and UCSF Chimera. Among them, Benzaldehyde, 5-bromo-2-hydroxy-, (5-trifluoromethyl-2pyridyl)hydrazone exhibited the highest binding affinity (-8.2 kcal/mol), confirming its strong interactions at the active site of B175L. Molecular dynamics (MD) simulations demonstrated the compound’s stability, with root mean square deviation and fluctuation (RMSD, RMSF) indicating minimal conformational changes (<2 Å). The compound maintained stable hydrogen bonds and hydrophobic interactions, reinforcing structural robustness. Taken together, these results clearly emphasize *T. cordifolia-*derived Benzaldehyde, 5-bromo-2-hydroxy-, (5-trifluoromethyl-2-pyridyl)hydrazone as a promising B175L inhibitor, advancing exploration towards effective antiviral solutions for ASFV.

## Introduction

African swine fever (ASF) is a highly lethal and contagious viral disease that affects both domestic and wild pigs. Acute infections are often fatal, with a mortality rate approaching 100%. ASF is classified as a notifiable disease by the World Organization for Animal Health (WOAH) due to its severe economic and animal health impact (1). The disease was first documented in Kenya in 1921 and has since become a global concern. The current transboundary epidemic began with the introduction of the highly virulent ASF virus (ASFV) genotype II into Georgia in 2007 (2). As of November 2024, ASF has been reported in 63 countries and territories spanning five major regions. Between January 2022 and November 2024, the disease affected over 781,000 domestic pigs and 27,400 wild boars, leading to the loss of more than 1.87 million animals (3). ASFV belongs to the genus *Asfivirus* within the family *Asfarviridae*. It is a large, complex, double-stranded DNA virus with genome sizes ranging from 170 to 194 kilobases (kb) among different isolates. The viral genome encodes approximately 150 to 200 proteins involved in diverse functions, including viral structure formation, replication, and immune evasion (4).

The lack of commercially approved vaccines or effective therapeutics for African swine fever (ASF) has escalated into a major crisis for the global pork industry (5). Although live-attenuated vaccines (LAVs) have shown potential, their use is limited by safety concerns, including the risk of reversion to virulence and the possibility of persistent infections. Subunit vaccines, while generally safer, still require further investigation to identify effective antigens and suitable adjuvants capable of eliciting strong and durable immune responses (6). In addition to these scientific hurdles, vaccine development faces practical challenges such as adverse side effects, limited diagnostic capabilities, data insufficiency for risk assessments, biological complexity, and high research costs (7). As a result, there is an urgent need to explore alternative strategies for ASF control, particularly through the discovery of safe and effective antiviral agents (8). Natural plant-derived compounds offer a promising alternative for antiviral drug discovery due to their vast chemical diversity and established antiviral potential. Numerous medicinal plants have demonstrated efficacy in inhibiting viral infections, and extensive research supports the antiviral activity of plant-derived bioactive compounds in both *in vitro* and *in vivo* models (2). In this context, computer-aided drug design (CADD), also known as *in silico* drug discovery, has emerged as a powerful tool in modern pharmacological research. These computational approaches are not only cost-effective and time-efficient but also minimize the reliance on animal testing in early-stage drug development (9).

*Tinospora cordifolia* is a traditional medicinal plant with a long-standing history in Ayurvedic medicine and is renowned for its wide spectrum of pharmacological activities. It is widely distributed across tropical regions, particularly in India and Southeast Asia, and is also endemic to Sri Lanka. The plant is known for its immunomodulatory, anti-inflammatory, antiviral, antioxidant, and antimicrobial properties, which are primarily attributed to its rich array of bioactive compounds, including alkaloids, glycosides, diterpenoid lactones, and polysaccharides. A substantial body of literature supports its immunomodulatory effects and its demonstrated efficacy against viral infections, including COVID-19 (10). Among the many proteins encoded by ASFV, B175L has recently been identified as a key immunomodulatory factor. (11)) reported that B175L plays a critical role in suppressing the type I interferon (IFN) response, a central antiviral defense mechanism. Specifically, B175L inhibits IFN-β production and downstream signaling by directly targeting STING (Stimulator of Interferon Genes) and its ligand 2′3′-cyclic GMP-AMP (2′3′-cGAMP), both essential components of the cyclic GMP-AMP synthase-STING (cGAS-STING) pathway. This interaction disrupts the activation of transcription factors IRF3 and NF-κB, ultimately suppressing IFN-stimulated gene expression. Due to its strong immune-suppressive function, B175L represents a promising therapeutic target for antiviral intervention.

Given that the ASFV B175L protein functions as a key negative regulator of the host’s antiviral interferon response through its interaction with STING, this *in silico* study aims to identify ligands derived from *T. cordifolia* that can effectively bind to B175L. By disrupting the B175L–STING interaction, the study seeks to restore STING-mediated innate immune signaling. This approach presents a novel and promising strategy for developing antiviral therapeutics against ASF.

## Materials and methods

### Collection, authentication, and preparation of plant material

In this study, the stems of *Tinospora cordifolia* were systematically collected in December 2024 from a privately owned herb garden placed in the Ratnapura district, Sri Lanka, and identified and authenticated by a Botanist (12) (13). Upon arrival at the laboratory, we thoroughly cleaned the herb stems to remove dirt or debris. The stems were cut into small pieces, sun-dried, and powdered using a mixer grinder. The powder obtained was sieved (0.2 mm) using a stainless-steel sieve and stored in an airtight polythene bag at -20 °C for further investigation (14).

### Plant Extraction Preparation

We weighed 7.5 g of the sample for extraction using an analytical balance. The sample was mixed with 100 mL of ≥99.9% methanol (VWR CHEMICALS; 85681.320) using a magnetic stirrer for 30 minutes at 40 °C. The mixture was filtered with Whatman No. 1 paper. Then the methanol was evaporated using a 30 °C water bath (Thermo Scientific, Precision Water Bath). The resulting pulp-like residual was used for further analysis. 15 mg of the residual was weighed into a 2 mL Eppendorf tube and dissolved with 1 mL of methanol for GC-MS analysis.

### Gas chromatography-mass spectrometry (GC-MS) analysis

GC-MS analysis of *T. cordifolia* stem methanolic extract was performed using an Agilent Technologies 7890B GC system with a 5977A mass selective detector and a 7890B autosampler. An Agilent DB FATWAX UI column (30 m × 0.250 mm × 0.25 µm) separated samples with helium carrier gas at 1.5 mL/min constant flow. Sample analysis occurred through 1 µL split mode injections (2:1) into an inlet set at 250 °C. The temperature protocol was started at 40 °C before incrementing to 115 °C at 5 °C/minute for 5 minutes, then to 140 °C at 5 °C/minute (5-minute hold), followed by a 2 °C/min increase to 210 °C that lasted 11 minutes and culminated with a 5 °C/minute increase to 240 °C maintained for 9 minutes. The complete assessment was run for 91 minutes. After sample preparation involving PTFE filtration, the direct injection was performed. The identification process was employed with the NIST14 and W9N11 libraries (15).

### Drug-likeness prediction

To evaluate the *in silico* ADME/T (absorption, distribution, metabolism, excretion, and toxicity) properties of the plant compounds, we used SwissADME (http://www.swissadme.ch/) and DataWarrior. SwissADME is an online tool that predicts drug-likeness and pharmacokinetic properties (16). The SMILES of the compounds were obtained from the PubChem database and inserted into the SwissADME interface (17). ADME properties were evaluated based on Lipinski’s rule of five, which suggests that compounds with more than 5 H-bond donors, 10 H-bond acceptors, a molecular weight over 500 Da, and a calculated Log P (MLogP) greater than 5 likely have poor absorption or permeation (18) (19). Mutagenicity, irritant, tumorigenicity, and reproductive toxicity-like parameters were listed as expected toxicities and evaluated using DataWarrior software (20) (21).

### Retrieval and characterization of protein sequences

The protein sequence for the uncharacterized protein B175L from the ASFV Georgia strain was obtained from the UniProt database (UniProt ID: A0A2X0RVG) (https://www.uniprot.org/uniprotkb?query=A0A2X0RVG) (22) (23). Chemical and physical attributes such as molecular weight, amino acid composition, theoretical pI, instability index, extinction coefficient, atomic composition, estimated half-life, the total number of positively charged residues (Arg + Lys), the total number of negatively charged residues (Asp + Glu), aliphatic index, and grand average of hydropathic were analyzed using ExPASy’s ProtParam tool (https://www.expasy.org/) (24).

### Three-dimensional (3D) structure prediction, refinement, and validation

The 3D structure of the protein was predicted using AlphaFold3 (https://alphafoldserver.com/), an advanced deep-learning-based protein structure prediction tool (25). The prediction was performed by feeding the primary amino acid sequence into the AlphaFold server. The quality of the predicted homology model was assessed using structural confidence metrics, including the predicted local distance difference test (pLDDT), predicted aligned error (PAE), predicted template modelling (PTM) score, and interchain predicted TM-score (ipTM) (26). The modeled protein 3D structure was refined using GalaxyWeb (https://galaxy.seoklab.org/). In homology modeling, structure validation is a vital step. The basic conformation of the suggested protein model was obtained by submitting it to ProSA-web (https://prosa.services.came.sbg.ac.at/prosa.php) (27).

### Assessment of model quality

The predicted 3D structure was evaluated using the SAVES 6.0 (https://saves.mbi.ucla.edu/) structure evaluation server, utilizing three assessments: PROCHECK, Verify3D, and ERRAT. PROCHECK visualized the backbone dihedral angles ψ against φ of amino acids in the protein structure (28) (29). Verify3D assessed the compatibility of a 3D atomic model with its corresponding 1D amino acid sequence (27). To further validate the protein model, MolProbity (https://molprobity.biochem.duke.edu/) was used, which provides a log-weighted score combining the clash score, percentage of Ramachandran outliers, and percentage of unfavorable side-chain rotamers (30).

### Molecular docking and visualization

For molecular docking, the 3D SDF conformation of all ADME/T-approved plant compounds was downloaded from PubChem (31) (32). The 3D SDF structures of compounds not available in PubChem were generated using ChemSketch software (Advanced Chemistry Development, Inc.) (33). Virtual screening docking was performed using the Vina Wizard function of PyRx software version 0.8, and Biovia Discovery Studio version 4.5 was used for docking pose visualization. The preparation of the protein involved converting pdb files into pdbqt format.

The docking procedure used a predefined grid box matching the predicted active site location. The CB-Dock2 server was employed for further docking evaluation (34), enabling the identification of interacting amino acids of respective CurPockets. Binding affinities were further confirmed using UCSF Chimera with the coordinates given by the CB-Dock server for grid center and box dimensions (35) (36). Biovia Discovery Studio was used for complex visualization (37) (38). The zinc finger binding area of B175L was identified using the Conserved Domain Database of NCBI (39).

### Molecular dynamics (MD) simulations

The MD simulations regarding the highest-ranked protein-ligand complex were performed using the Desmond module of the Schrödinger suite (40). The system was prepared through the System Builder tool, taking advantage of the simple point charge (SPC) water model as a solvent inside an orthorhombic box with periodic boundary conditions (PBC). To neutralize the system, the counter ions, Na+ and Cl-, were added. The OPLS4 force field was utilized to realize energy minimization and then MD simulations. The trajectory of the simulation spans 100.102 ns to record at 100 ps intervals under NPT ensemble conditions at a fixed pressure of 1.01 bar and temperature of 310 K. The total charge of the protein during the simulation was -5. Post-simulation analyses included analysis of root mean square deviation (RMSD) concerning protein and ligand, root mean square fluctuation (RMSF) to characterize local changes in protein and ligand, detailed protein-ligand contact analysis (including hydrogen bonds, hydrophobic, ionic, and water bridge interactions), second structure element (SSE) analysis and prediction of ligand properties such as radius of gyration (rGyr), intramolecular hydrogen bonds, molecular surface area (MolSA), solvent accessible surface area (SASA) and polar surface area (PSA) (41).

## Results

### Disease background

ASFV exhibits complex transmission dynamics involving both mechanical and biological routes, complicating prevention efforts. As shown in Fig 1A, mechanical transmission includes contaminated feed, vehicles, meat, flies, and personal contact, while biological transmission involves infected pigs and soft ticks. Understanding these different transmission routes is crucial for implementing effective biosecurity measures. Upon infection, pigs manifest a variety of clinical signs that include vomiting, diarrhea (sometimes with blood), internal bleeding, weakness, high fever, skin reddening, loss of appetite, abortion, and inability to stand (Fig 1B), resulting in a high mortality rate during acute cases. Rapidly detecting the clinical signs leads to a quick identification and diagnosis of ASF outbreaks. The spread of the disease has been observed across various continents, with new outbreaks reported in numerous countries at an increasingly rapid rate (Fig 1C). The large double-stranded DNA genome of ASFV further exemplifies its complex nature, encoding a multitude of viral proteins that contribute to immune evasion and pathogenesis, impairing the development of effective treatments (Fig 1D).

**Fig 1.**
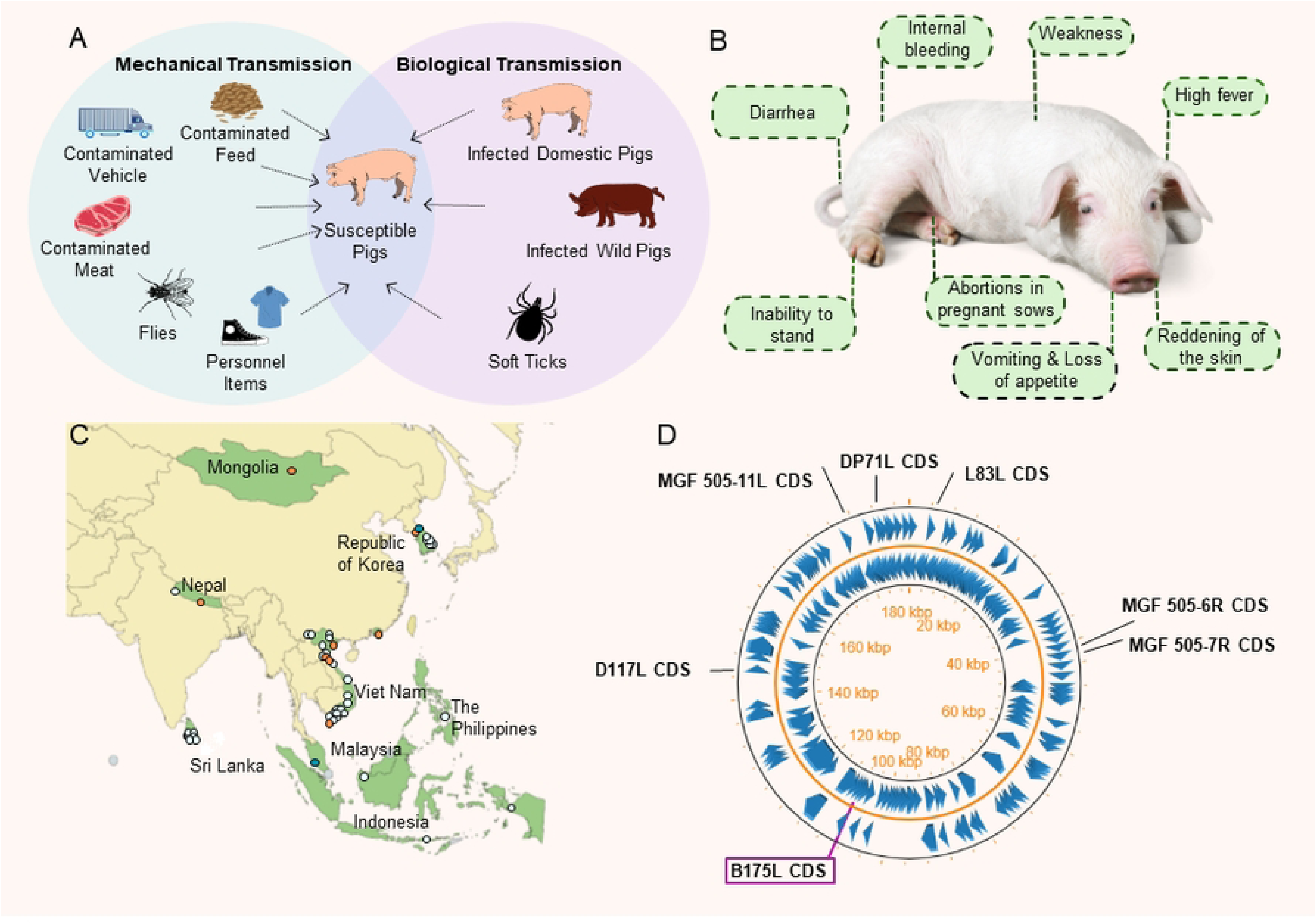
Overview of African swine fever (ASF) (A) Transmission pathways of ASF virus (ASFV) (42). (B) Key symptoms of ASF. (C) ASF situation in Asia for the last ten weeks up to 20^th^ February 2025. New outbreaks reported appear in orange, and new detections in free wild boar appear in dark blue from 6/02/2025 to 20/02/2025. Detection in the farm or free wild boar during 12/12/2024-6/02/2025 appears in light blue. Source: FAO, 2025. Map of the ASF situation in Asia. (Cited March 2025). (D) Circular representation of the ASFV genome. The ASFV Georgia 2007/1 isolate genome is shown. This visualization clearly emphasizes the location of various coding sequences (CDS). Among the 195 proteins encoded by the genome, this visualization clearly emphasizes MGF 505-7R, MGF 505-11L, L83L, D117L, MGF 505-6R, DP71L, and B175L proteins, which have been identified as STING-mediated inhibitors. The B175L protein specifically highlighted in the figure exhibits a length of 528 bp CDS located on the reverse strand between positions 108, 527 bp, and 109,054 bp.

### Plant description and research workflow

The *in silico* methodology presents a promising approach for identifying novel inhibitors against ASFV, derived from plants. Understanding the taxonomic lineage (Fig 2A) of *T. cordifolia* (Kingdom-Plantae, Phylum-Tracheophyta, Class-Magnoliopsida, Order-Ranunculales, Family-Menispermaceae, Genus-*Tinospora*, Species-*Tinospora cordifolia*) within the plant kingdom is essential for recognizing its potential inhibiting compounds for the virus. As shown in Fig 2B, *T. cordifolia* exhibits diverse pharmacological properties such as anti-diabetic, anti-oxidant, antitoxic, immunomodulatory, anti-microbial, anti-inflammatory, anti-HIV, and anti-cancer, showcasing its medicinal significance, making it a possible candidate for inhibiting ASFV. The *in silico* workflow followed in this study, including plant selection, extraction preparation, GC-MS analysis, ADME/T analysis, and in parallel to that, homology modeling, binding site identification, molecular docking, and MD simulations (Fig 2C), uncovers novel inhibitors against ASFV, facilitating more effective and efficient methodology and results.

**Fig 2.**
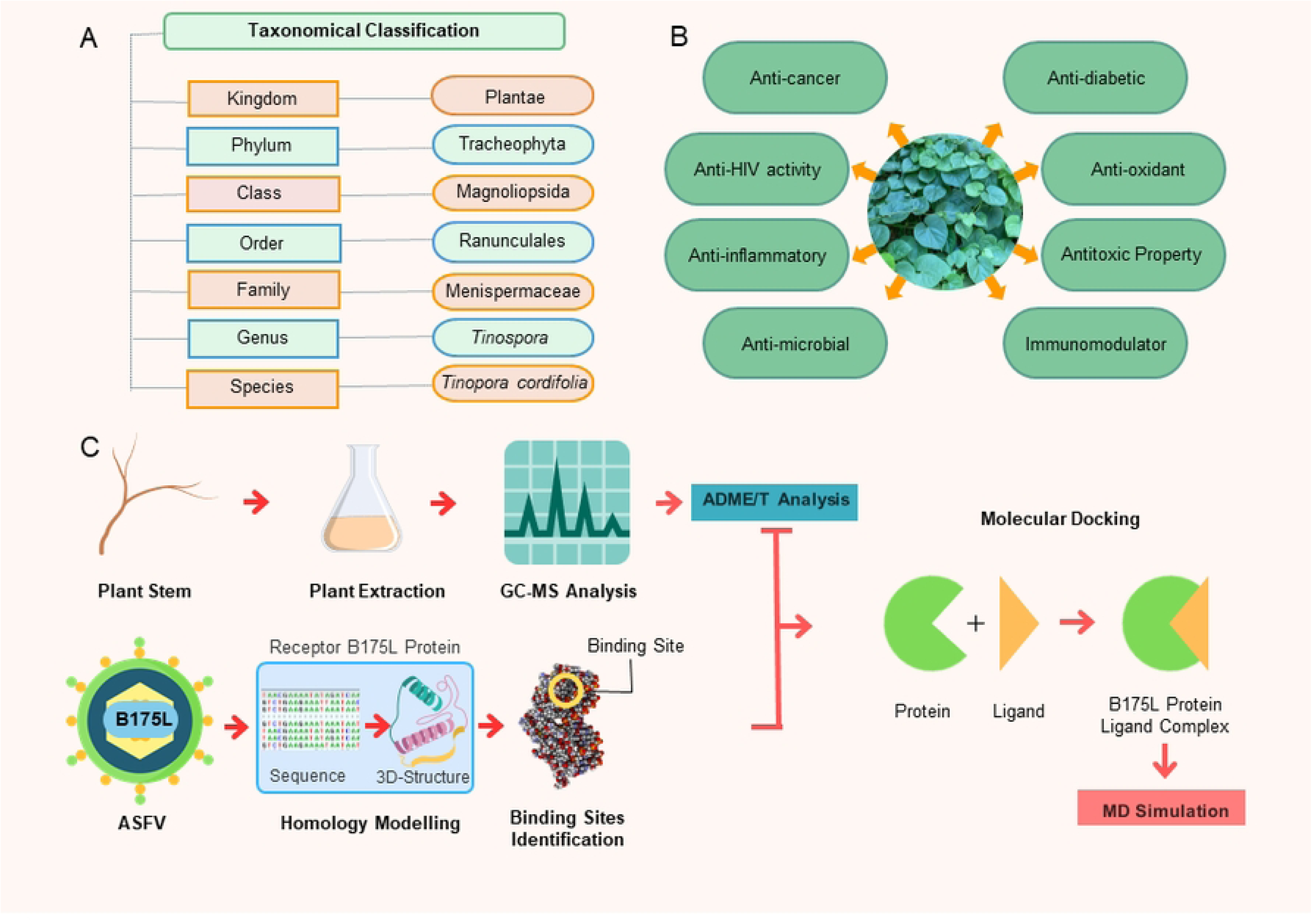
Comprehensive illustration of *Tinospora cordifolia* and study workflow. (A) Taxonomy of *T. cordifolia*. (B) Pharmacological properties of *T. cordifolia* (43). (C) Computational screening workflow for identifying B175L inhibitors in *T. cordifolia*.

### Plant stem extract profiling via GC-MS

GC-MS analysis of the methanolic extract of *T. cordifolia* stems recorded a total of 92 peaks (S1A Fig) corresponding to bioactive compounds. These compounds were identified by relating their peak retention time, peak area (%), height (%), and mass spectral fragmentation patterns to known compounds. The results revealed that 124 compounds were identified in the methanol extract. The phytoconstituents in the methanol stem extract of *T. cordifolia* included butanoic acid derivatives, aromatic aldehydes (e.g., benzaldehyde), and fatty acid esters (e.g., hexadecanoic acid methyl ester) etc. Key compounds identified were butanoic acid, 4,4,4-trichloro-2-methyl-, methyl ester, 1,1,2-trichloro-2-[(2-chloroethyl)thio]ethene, benzaldehyde, 5-bromo-2-hydroxy-,(5-trifluoromethyl-2-pyridyl)hydrazone, methanamine, N-methoxy, methanamine, N-hydroxy-N-methyl-, 1,6-heptadiyne, 4-benzyloxy-3-methoxyphenylacetonitrile, carbamic acid, N-(3-oxo-4-isoxazolidinyl)-, benzyl ester, butanal, 3-methyl-, pentanal, hydrazine, 1,2-dimethyl-, acetic acid, hydrazine, 1,1-dimethyl-, 2,4-dihydroxy-2,5-dimethyl-3(2H)-furan-3-one, hydrazine, 1,2-dibutyl, butanoic acid, 4-hydroxy-, butyrolactone, 4-ethylbenzoic acid, 2-butyl ester, oxime-, methoxy-phenyl-_, phthalic acid, ethyl tetradecyl ester, 2-furanmethanol, 4-ethylbenzoyl chloride, 1,2,4-benzenetricarboxylic acid, cyclic 1,2-anhydride, nonyl ester, 1H-indol-5-ol, oxalic acid, cyclohexylmethyl tridecyl ester, 3-cyclohexen-1-ol, 3-methyl-, succinic acid, dodec-2-en-1-yl cis-4-methylcyclohexyl ester, heptane, 1,7-dibromo-, 1,2-cyclopentanedione, 2-cyclopenten-1-one, 2-hydroxy-, 3H-pyrazol-3-one, 1,2-dihydro-5-methyl-, mequinol, phenol, 2-methoxy-, benzyl nitrile, cyclododecane, cyclopropane, nonyl-, 1-tetradecanol, docosane, eicosyl isobutyl ether, hexadecanoic acid, methyl ester, pentadecanoic acid, 14-methyl-, methyl ester, 4H-pyran-4-one, 2,3-dihydro-3,5-dihydroxy-6-methyl-, 4H-pyran-4-one, 3,5-dihydroxy-2-methyl-, sarcosine anhydride, glycerin, 3-methoxy-2,2-dimethyloxirane, butanedioic acid, monomethyl ester, benzofuran, 2,3-dihydro-, benzene, 1-ethynyl-4-fluoro-, cumidine, tetracosane, cis-13-octadecenoic acid, methyl ester, 9-octadecenoic acid, methyl ester, (E)-, 9,12-octadecadienoic acid (Z,Z)-, linoelaidic acid, tricyclo[6.3.0.0(2,6)]undecan-10-one, 3-[(2-methoxyethoxy)methoxy]-2-methyl-, 3,3’-isopropylidenebis(1,5,8,11-tetraoxacyclotridecane), 2-[2-[2-[2-[2-[2-[2-[2-(2-hydroxyethoxy)ethoxy]ethoxy]ethoxy]ethoxy]ethoxy]ethoxy]ethoxy]ethanol, methyl 3-hydroxybutyrate, TMS derivative, 2-[2-[2-[2-[2-[2-[2-[2-[2-(2-hydroxyethoxy)ethoxy]ethoxy]ethoxy]ethoxy]ethoxy]ethoxy]ethoxy]ethoxy]ethanol, 9,12,15-octadecatrienoic acid, (Z,Z,Z)-, methyl 8,11,14-heptadecatrienoate, decaethylene glycol, TMS derivative, undecaethylene glycol, TMS derivative, 18,18’-bi-1,4,7,10,13,16-hexaoxacyclononadecane, 2-[2-[2-[2-[2-[2-[2-(2-hydroxyethoxy)ethoxy]ethoxy]ethoxy]ethoxy]ethoxy]ethoxy]ethanol, 15-crown-5, 1,4,7,10,13,16-hexaoxacyclooctadecane, heptaethylene glycol, 2-[2-[2-[2-[2-[2-[2-(2-methoxyethoxy)ethoxy]ethoxy]ethoxy]ethoxy]ethoxy]ethoxy]ethanol, 2-[2-[2-[2-[2-[2-[2-[2-(2-methoxyethoxy)ethoxy]ethoxy]ethoxy]ethoxy]ethoxy]ethoxy]ethoxy]ethanol, 2-[2-[2-[2- [2-[2-[2-[2-(2-hydroxyethoxy)ethoxy]ethoxy]ethoxy]ethoxy]ethoxy]ethoxy]ethoxy]ethanol, hexaethylene glycol, 2-[2-[2-[2-[2-[2-[2-[2-(2-methoxyethoxy)ethoxy]ethoxy]ethoxy]ethoxy]ethoxy]ethoxy]ethoxy]ethanol, (18S,19S)-18,19-dihydroxy-1,4,7,10,13,16-hexaoxocycloencosane, 3-(1,3-dihydroxyisopropyl)-1,5,8,11,14,17-hexaoxacyclononadecane, 3,6,9,12,15-pentaoxanonadecan-1-ol.

### *In silico* ADME and toxicity prediction

Initial GC-MS analysis identified 124 phytochemical compounds as potential therapeutic candidates. These underwent ADME evaluation using SwissADME, strictly adhering to Lipinski’s Rule of Five. 119 compounds (95.97%) satisfied all criteria, demonstrating favorable pharmacokinetic properties for oral bioavailability. Five compounds were excluded due to violations of more than one rule. ADME-compliant compounds were further analyzed for toxicity endpoints using DataWarrior software. As shown in Table 1, 52 compounds (43.70%) exhibited no predicted toxicity flags across all categories, focusing on four critical parameters: mutagenicity, tumorigenicity, reproductive toxicity, and irritant potential.

**Table 1.**
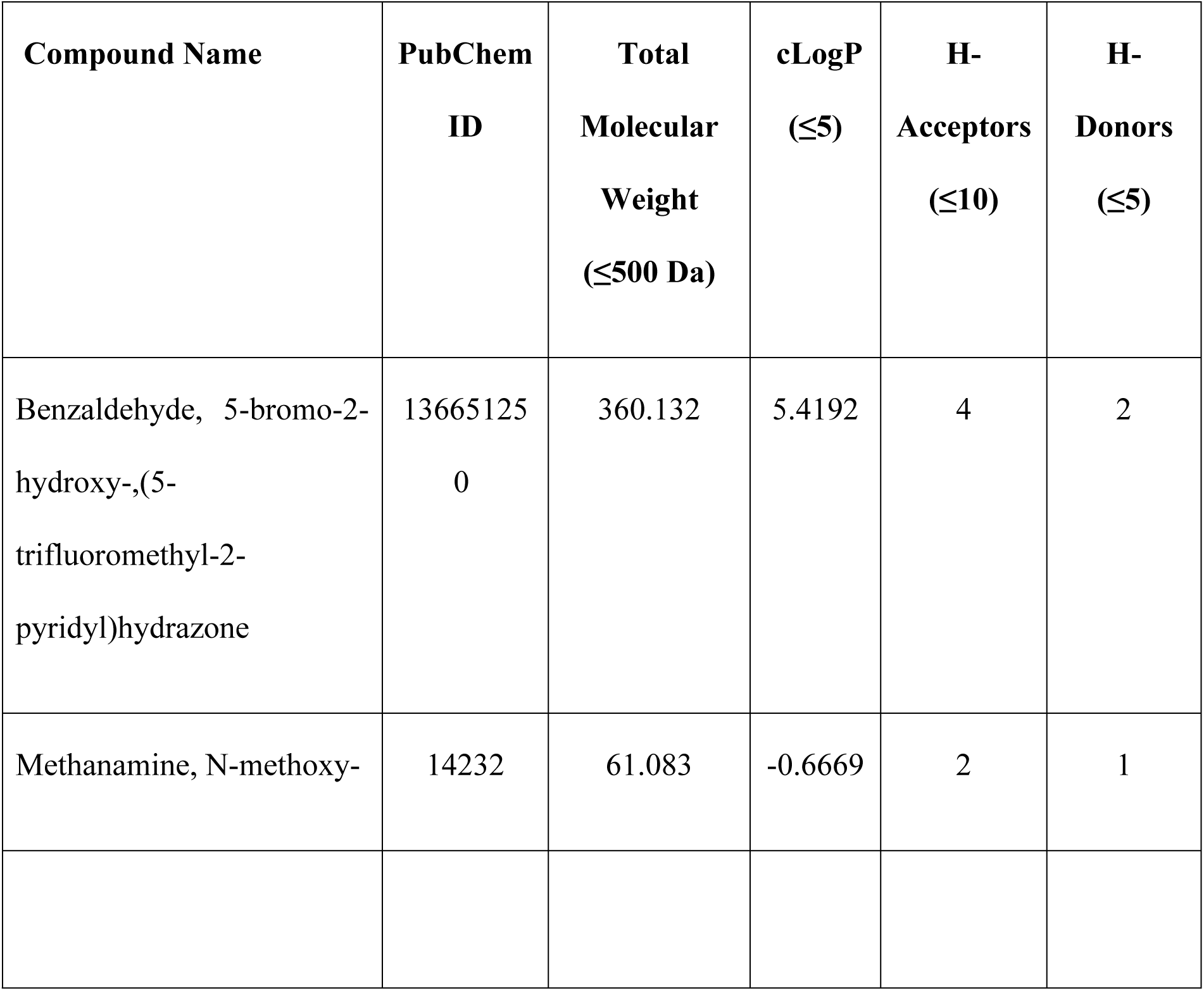
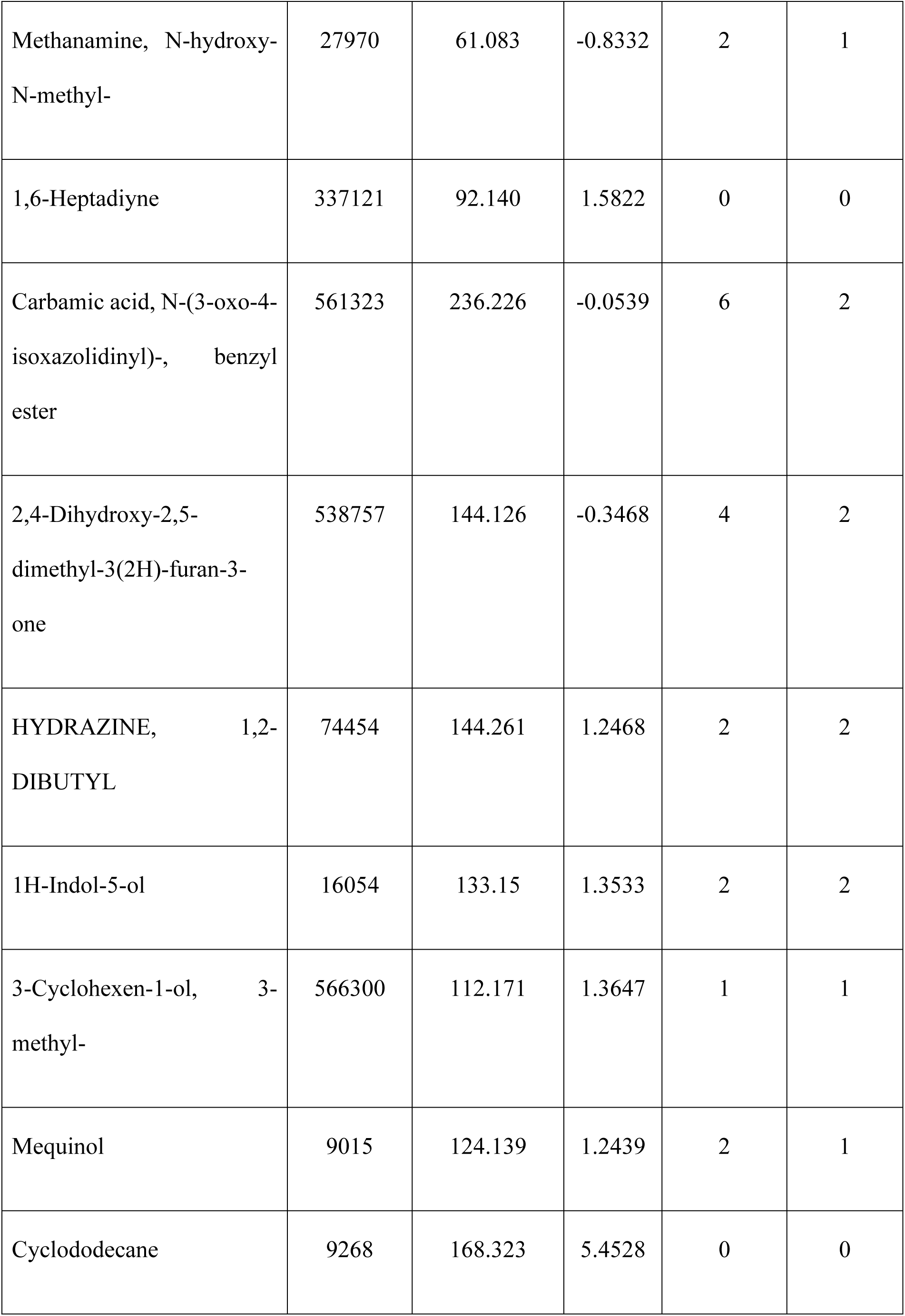
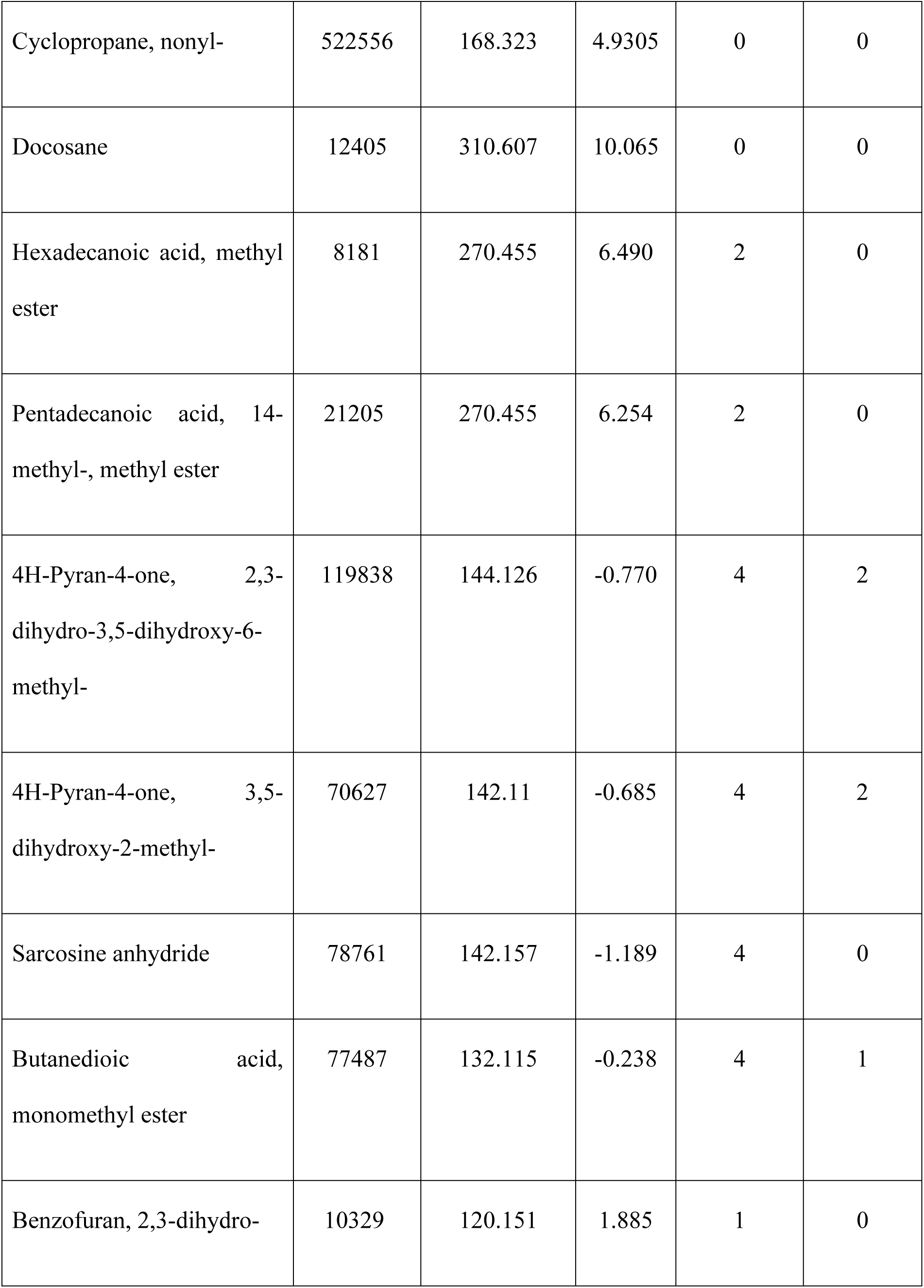
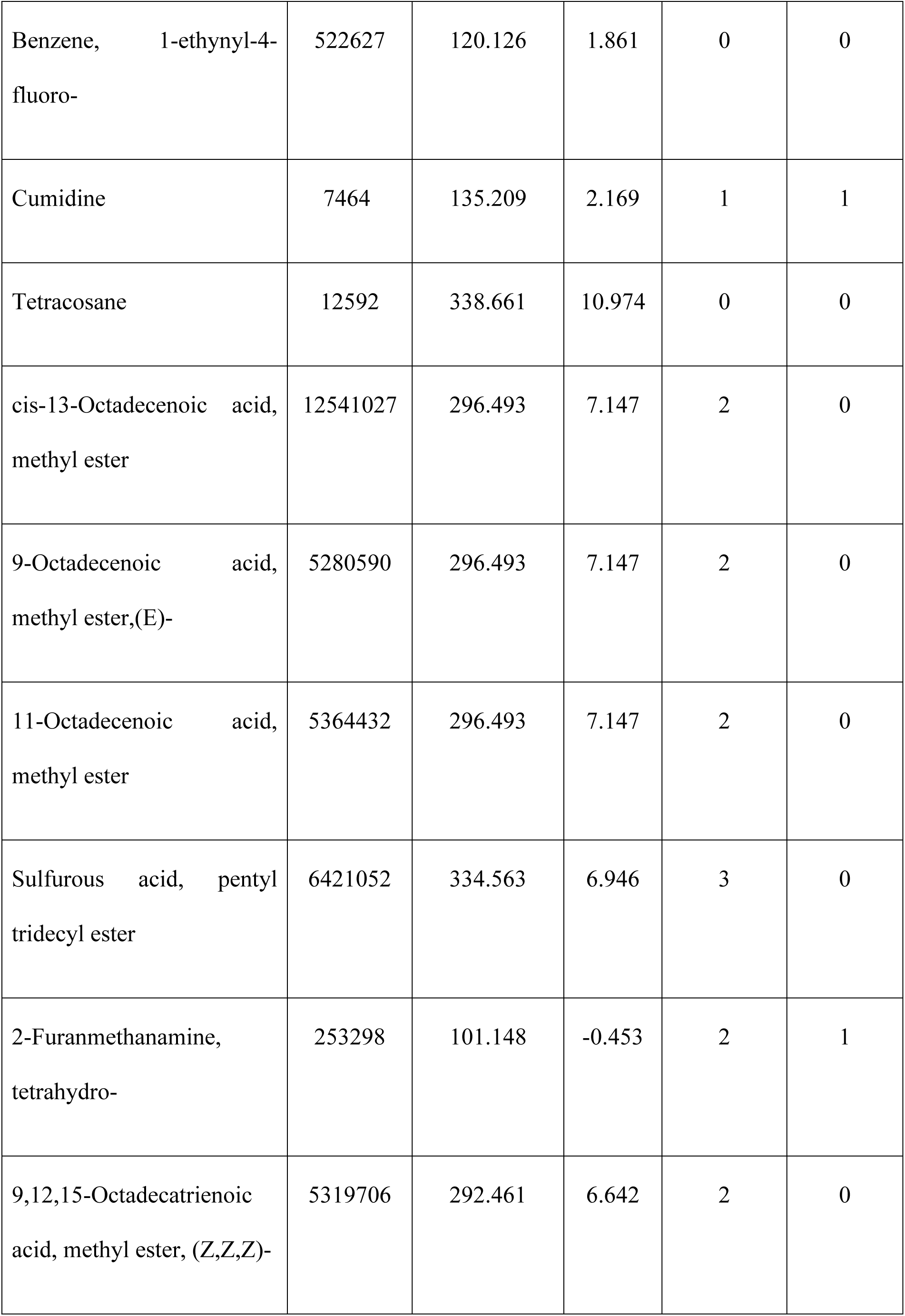
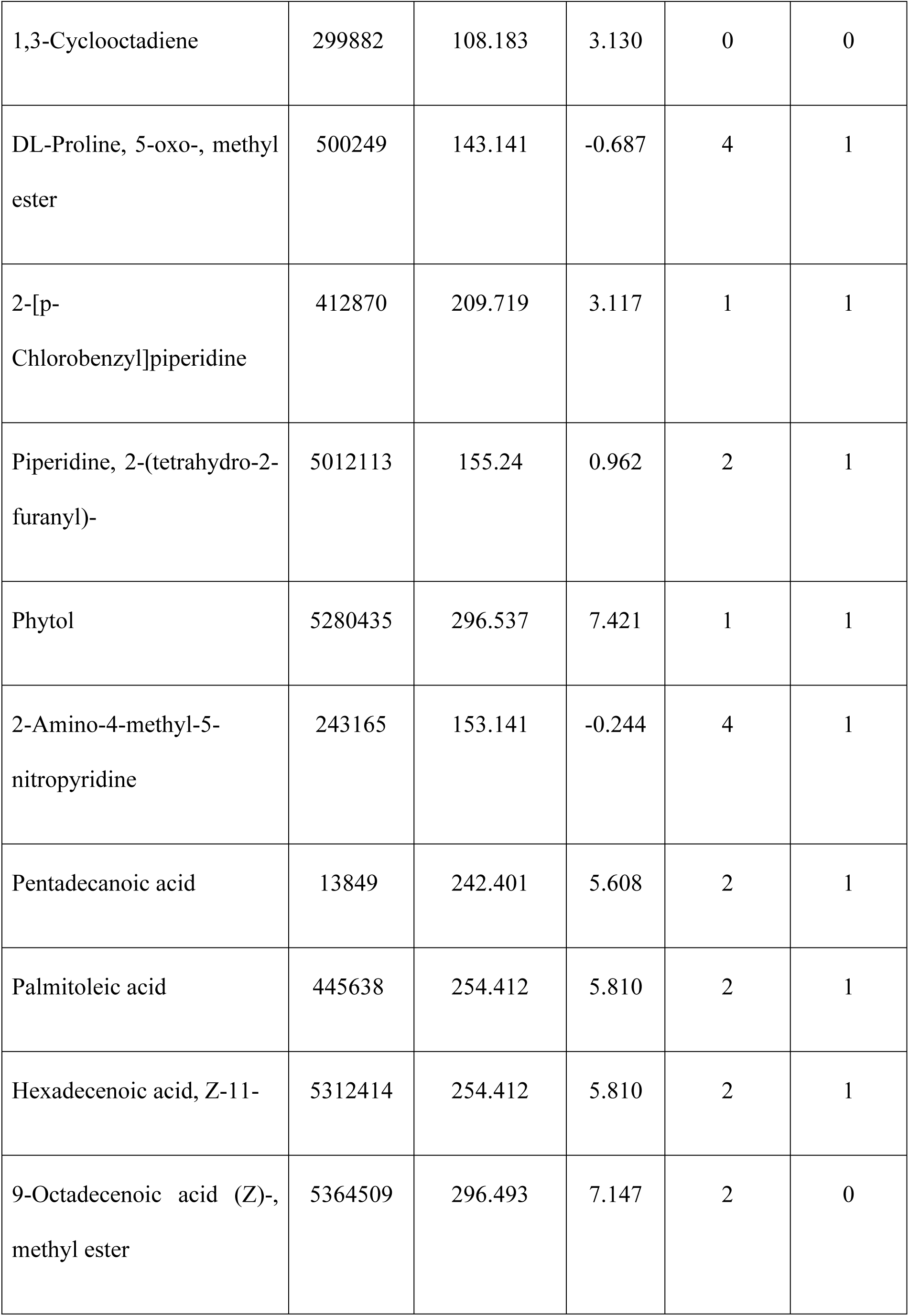
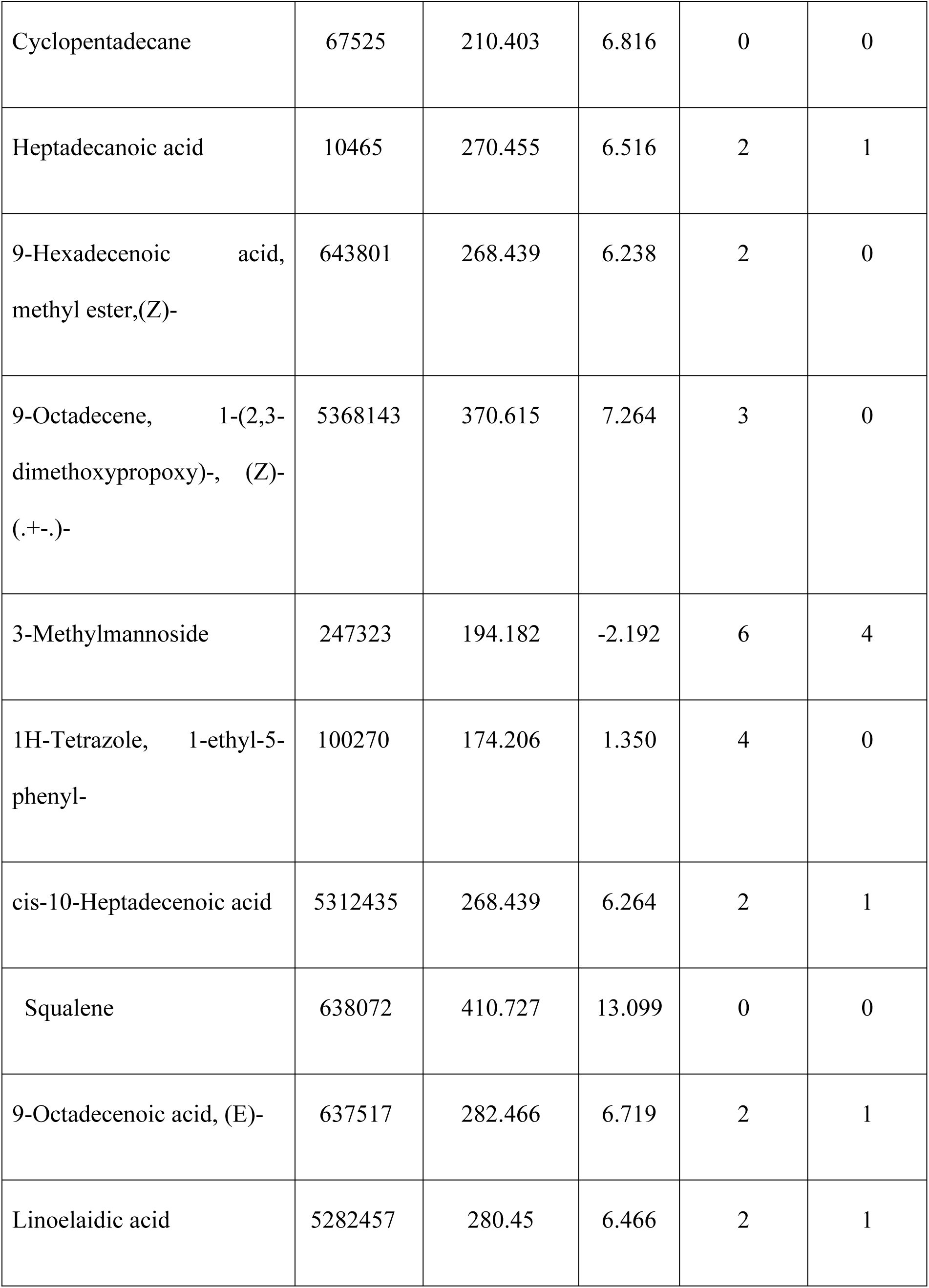
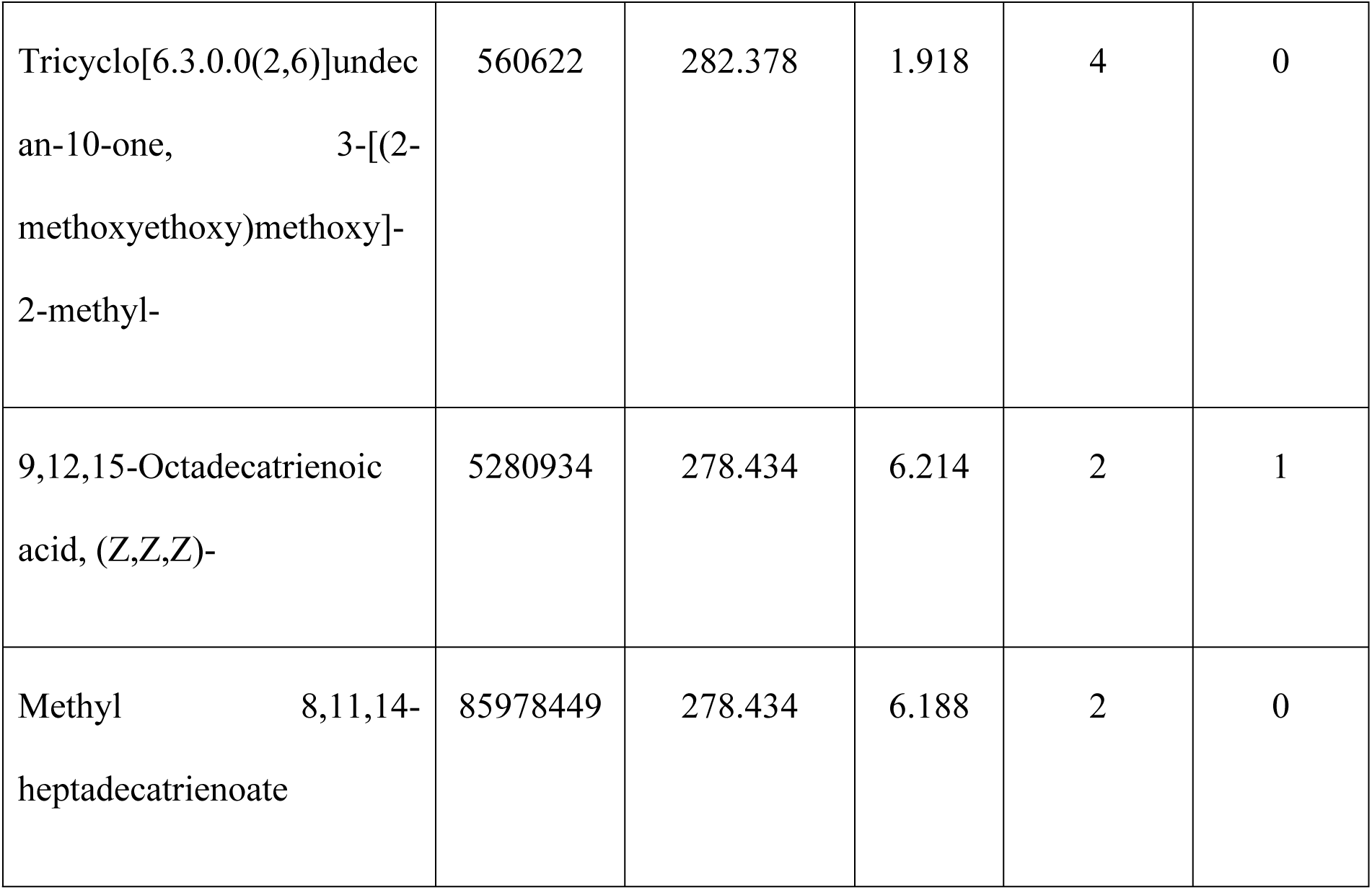
ADME properties of non-toxic compounds from *Tinospora cordifolia*.

### Physicochemical analysis of the B175L

As shown in Fig 3A the protein consists of 175 amino acids, among the most abundant was Glu (E) 19 followed by Ser (S) 17, Thr (T) 16, Leu (L) 13, Lys (K) 13, Ile (I) 10, Phe (F) 10, Tyr (Y) 9, Val (V) 9, Ala (A) 7, Asn (N) 7, Pro (P) 7, Arg (R) 6, Cys (C) 6, Gly (G) 6, His (H) 5, Met (M) 5, Asp (D) 4, Trp (W) 4, Gln (Q) 2, Pyl (O) 0, and Sec (U) 0. The compound molecular weight was 20348.20 Da, with a theoretical isoelectric point of 5.77, indicating a positively charged protein. The total number of negatively charged (Asp + Glu) residues and positively charged (Arg + Lys) residues was discovered to be 23 and 19, respectively. The protein was classified as unstable by the computed instability index (II) of 52.85. The aliphatic index was 70.17. The Grand Average of Hydropathy (GRAVY) value was -0.355. Mammalian reticulocytes (*in vitro*) were found to have a half-life of 30 hours, > 20 hours for yeast, and >10 hours for Escherichia coli (Fig 3B). The molecular formula of protein was identified as C_917_H_1395_N_229_O_273_S_11_ with a total of 2825 atoms (Fig 3C).

**Fig 3.**
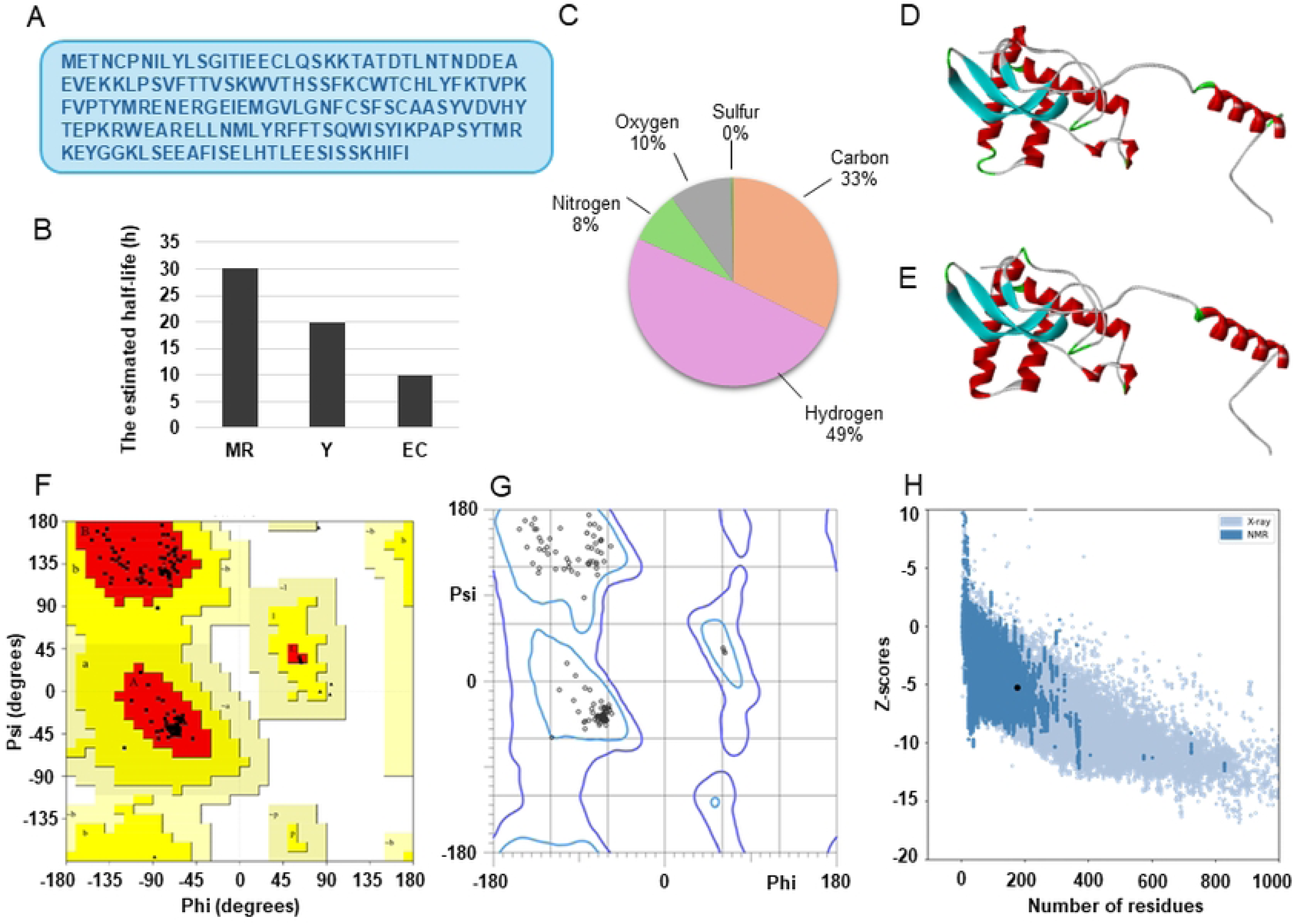
Physicochemical characterization and modelling of ASFV B175L. (A) Amino acid sequence of B175L. (B) Predicted half-life of B175L in mammalian reticulocytes (MR), yeast (Y), and *Escherichia coli* (EC) (a, b, and c figures were generated based on details obtained from the ProtParam tool on the ExPASy server). (C) Atomic composition of B175L. (D) AlphaFold3 modeled structure of B175L visualized using Discovery Studio. (E) GalaxyWeb refined the structure of B175L, visualized using Discovery Studio. Ramachandran plot of the homology modeled and refined of B175L obtained from (F) PROCHECK and **(G)** MolProbity. (H) ProSA z-core diagram.

### Confidence metrics and refinement outcomes of B175L

We generated a structural model of the ASF B175L protein using AlphaFold3 (S1B Fig). The model provides insights into the protein’s overall fold and predicted structural characteristics. AlphaFold3 confidence metrics were used to assess the reliability of the generated structure. The selected AlphaFold3 model exhibited a pTM of 0.72, indicating a generally high confidence level in the overall structural prediction. The ranking score, which reflects AlphaFold3’s internal assessment of model quality, is more than >0.80, suggesting that this model is among the higher-quality predictions for B175L. Analysis of the model revealed that a fraction of the protein (0.26) is predicted to be disordered. This suggests the presence of flexible regions within the B175L structure. The model exhibited a ‘has clash’ value of 0.0, indicating that no significant steric clashes were detected in the predicted conformation. As our analysis focused on the B175L protein in isolation, the model was generated without multiple chains. As a result, ipTM was null. The chain pair ipTM score was 0.72, and the chain pair pae min was 0.76. The projected protein’s tertiary structure was refined using the Galaxy web server, generating five refined models and increasing the number of amino acids in favored locations. Compared to the initial model, the best-refined model was selected based on MolProbity, Global Distance Test - High Accuracy (GDT/HA), Root Mean Square Deviation (RMSD), clash score values, and poor rotamers in the structures (For model 4 - MolProbity value=1.256, GDT-HA=0.9571, RMSD=0.416, clash score=4.9, poor rotamers=0.6). The AlphaFold3 model and refined model 4 were chosen and visualized in Discovery Studio (Fig. 3D and E).

### Quality assessment of the predicted model

The Ramachandran plot was obtained from two web servers, PROCHECK and MolProbity, and the results are summarized in Table 2 PROCHECK evaluated the scalability of the galaxy server refined model through a Ramachandran plot analysis. The VERIFY3D score of the modeled protein indicated that 67.43% of the protein residues have an average 3D-1D score ≥0.1. The ERRAT quality factor was found to be 87.6623. The Ramachandran plot analysis revealed no residue within disallowed regions (Figs 3F and G). In addition, ProSA web server analysis resulted in a Z score of -5.26, indicating the model validation (Fig 3H).

**Table 2.** Ramachandran quality parameter for homology model using PROCHECK and MolProbity servers.

| Servers | Residues in<br>allowed region (%) | Residues in<br>outlier region (%) |
| --- | --- | --- |
| PROCHECK | 100% | 0 |
| MolProbity | 100% | 0 |

### Computational docking and structural visualization

The 52 ADME/T-compliant ligands underwent virtual screening against the refined B175L protein model of AFSV using PyRx 0.8 (AutoDock Vina). A binding affinity cutoff of more than -6 kcal/mol identified compounds, with affinities distributed as follows: 1 ligand at -8.5 kcal/mol, 3 ligands between −6.1 and −7.0 kcal/mol. The top candidate exhibited a binding affinity of −8.5 kcal/mol (Fig 4A). The top 4 candidates advanced to CB-Dock2 validation (Fig 4B), confirming robust target engagement. Subsequent binding mode verification of the highest binding affinity having ligand was performed in UCSF Chimera (grid center: x = 10, y = 1, z = 8; box dimensions: 22 × 22 × 22 Å³) and revealed stable interactions within the B175L active site by having a binding affinity of -8.2 kcal/mol. This multi-tiered computational strategy prioritized 1 lead compound, Benzaldehyde, 5-bromo-2-hydroxy-, (5-trifluoromethyl-2-pyridyl)hydrazone (L1), with superior binding characteristics for further investigation (Fig 4C).

**Fig 4.**
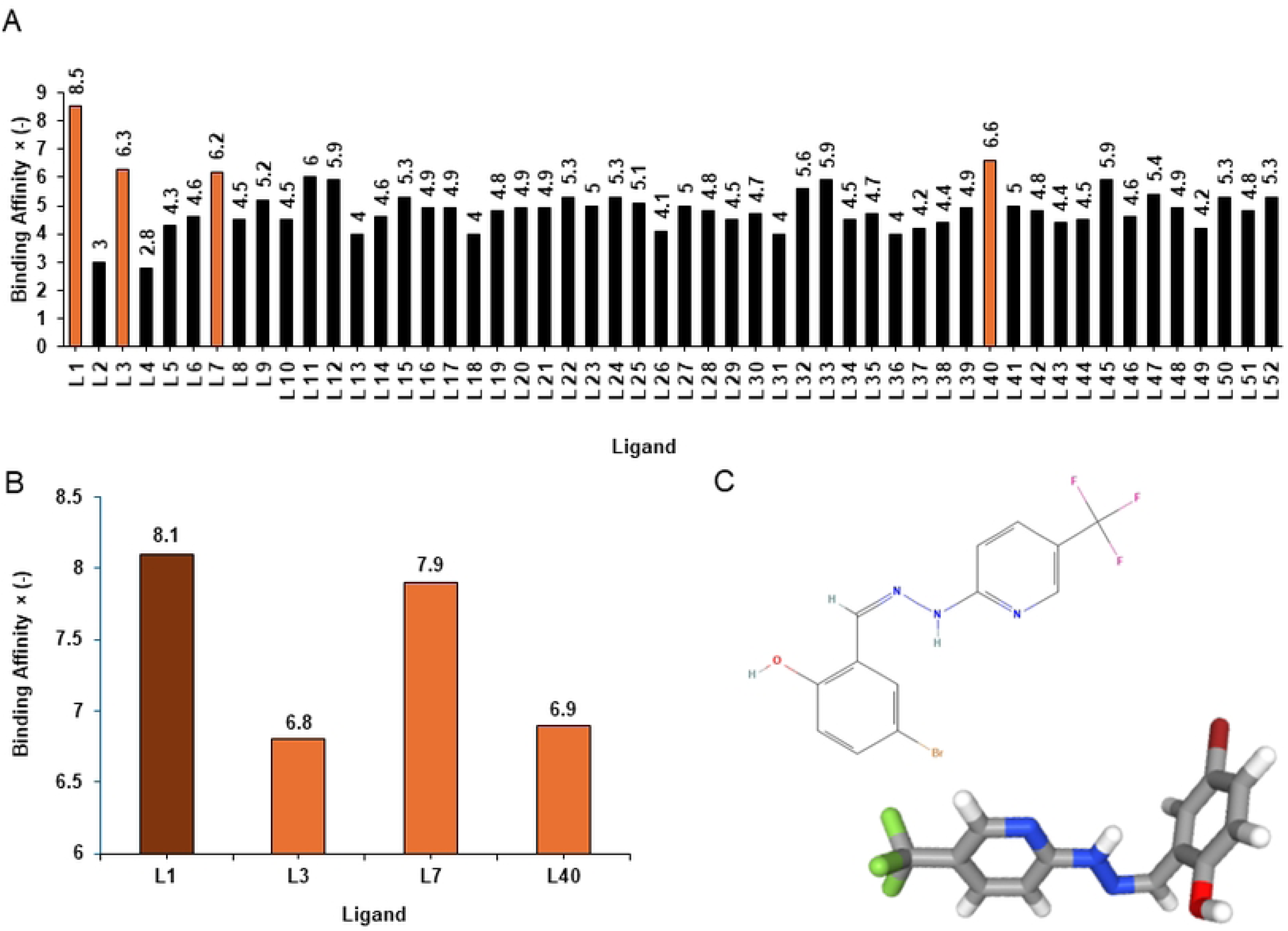
Graphical illustration of docking scores from PyRx virtual screening and CB-Dock2. (A) PyRx virtual screening results. This graph clearly illustrates docking scores for 52 ADME/T adhered ligands. Ligands with a binding affinity greater than -6 kcal/mol are represented by orange color bars. (B) CB-Dock2 molecular docking results for the top 4 ligands. The ligand with the highest binding affinity is indicated by a brown color bar. L1-Benzaldehyde, 5-bromo-2-hydroxy-,(5-trifluoromethyl-2-pyridyl)hydrazone, L2-Methanamine, N-methoxy-, L3-1H-Indol-5-ol, L4-Methanamine, N-hydroxy-N-methyl-, L5-1,6-Heptadiyne, L6-2,4-Dihydroxy-2,5-dimethyl-3(2H)-furan-3-one, L7-Carbamic acid, N-(3-oxo-4-isoxazolidinyl)-, benzyl ester, L8-3-Cyclohexen-1-ol, 3-methyl-, L9-HYDRAZINE, 1,2-DIBUTYL, L10-Mequinol, L11-Cyclododecane, L12-Cyclopropane, nonyl-, L13-Docosane, L14-Hexadecanoic acid, methyl ester, L15-Pentadecanoic acid, 14-methyl-, methyl ester, L16-4H-Pyran-4-one, 2,3-dihydro-3,5-dihydroxy-6-methyl-, L17-4H-Pyran-4-one, 3,5-dihydroxy-2-methyl, L18-Sarcosine anhydride, L19-Butanedioic acid, monomethyl ester, L20-Benzofuran, 2,3-dihydro-, L21-Benzene, 1-ethynyl-4-fluoro-, L22-Cumidine, L23-Tetracosane, L24-cis-13-Octadecenoic acid, methyl ester, L25-9-Octadecenoic acid, methyl ester,(E)-, L26-11-Octadecenoic acid, methyl ester, L27-11-Octadecenoic acid, methyl ester, L28-2-Furanmethanamine, tetrahydro-, L29-9,12,15-Octadecatrienoic acid, methyl ester, (Z,Z,Z)-, L30-1,3-Cyclooctadiene, L31-DL-Proline, 5-oxo-, methyl ester, L32-2-[p-Chlorobenzyl]piperidine, L33-Piperidine, 2-(tetrahydro-2-furanyl)-, L34-Phytol, L35-2-Amino-4-methyl-5-nitropyridine, L36-Pentadecanoic acid, L37-Palmitoleic acid, L38-Hexadecenoic acid, Z-11-, L39-9-Octadecenoic acid (Z)-, methyl ester, L40-Cyclopentadecane, L41-Heptadecanoic acid, L42-9-Hexadecenoic acid, methyl ester,(Z)-, L43-9-Octadecene, 1-(2,3-dimethoxypropoxy)-, (Z)-(.+-.)-, L44-3-Methylmannoside, L45-1H-Tetrazole, 1-ethyl-5-phenyl-, L46-cis-10-Heptadecenoic acid, L47-Squalene, L48-Octadecenoic acid, (E), L49-Linoelaidic acid, L50-Tricyclo[6.3.0.0(2,6)]undecan-10-one, 3- [(2-methoxyethoxy)methoxy]-2-methyl-, L51-9,12,15-Octadecatrienoic acid, (Z,Z,Z)-, L52-Methyl 8,11,14-heptadecatrienoate. (C) Two-dimensional (2D) and three-dimensional (3D) structures of L1 ligand from the PubChem database.

### Docked pose interactions

To elucidate complex stability, an in-depth analysis of molecular interactions was performed for the best-docked complex (Figs 5A, 5B). The docked complex of Benzaldehyde, 5-bromo-2-hydroxy-, (5-trifluoromethyl-2-pyridyl)hydrazone-B175L exhibited the formation of 2 conventional hydrogen bonds with THR A:48, VAL A:46, as depicted in Fig 5C. It is responsible for increasing the strength and stability of ligand-protein binding. Studies indicate that the higher the number of hydrogen bonds, the higher the interaction strength between the ligand and the target. In addition to hydrogen bonding, the complex exhibited ten hydrophobic interactions at residues LEU43, LYS52, VAL46, VAL54, VAL91, PHE47, PHE74, PRO44, and PRO72. These hydrophobic interactions enhance binding affinity by creating buried non-polar contacts and anchoring the ligand in hydrophobic regions of the protein. Furthermore, secondary interactions were observed, including Pi-Pi T-shaped interactions at PHE47 and Pi-Alkyl interactions at residues PHE74, PRO44, LYS52, PRO44, VAL54, and VAL91. These supplementary interactions play a crucial role in fine-tuning ligand orientation and complementing primary bonds, thereby contributing to the overall stability of the complex. Two halogen bonds were identified in this interaction (Table 3).

**Fig 5.**
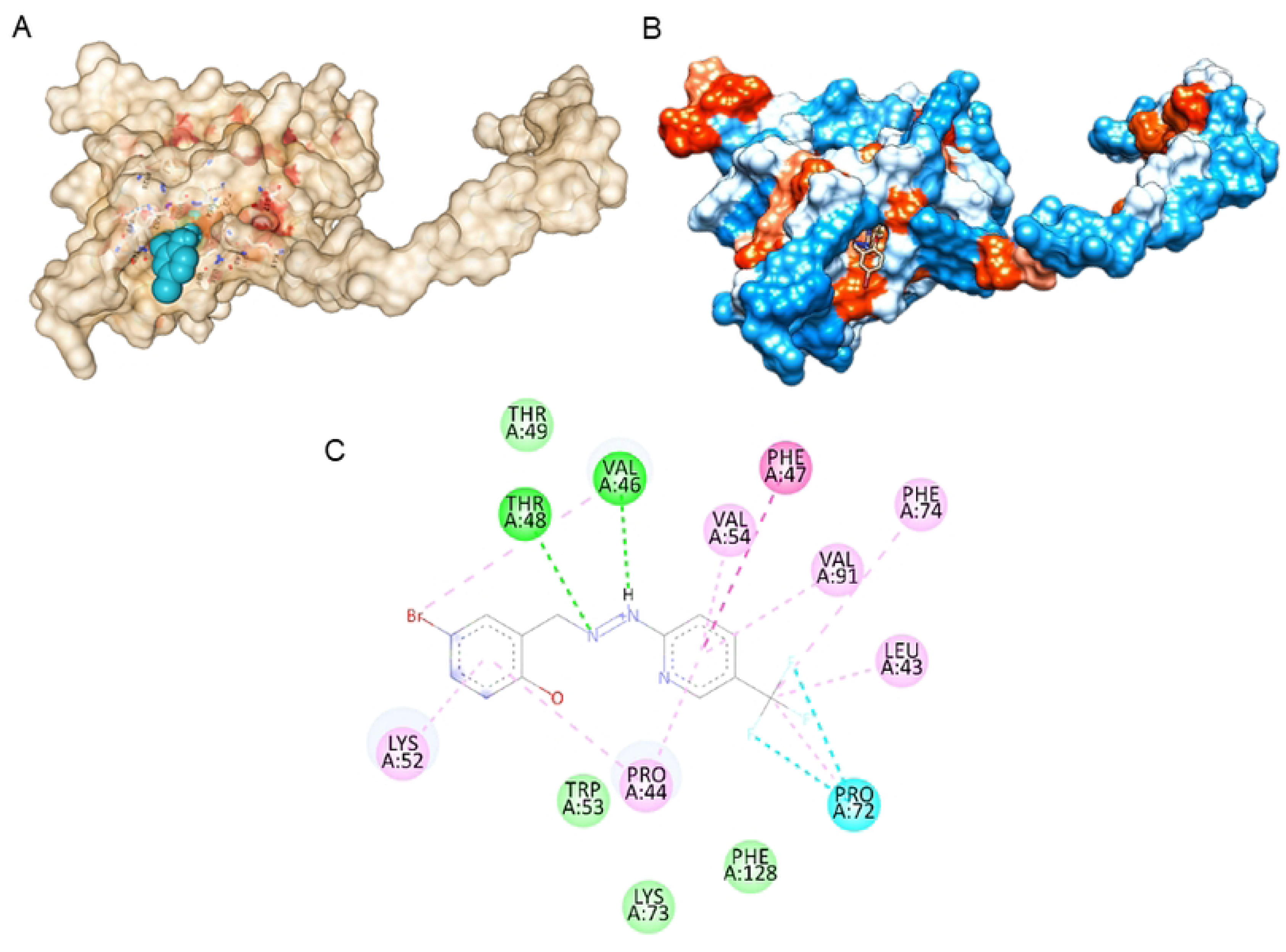
Protein-Ligand Complex Illustrations and Zinc Finger Binding Area of B175L. (A) Three-dimensional (3D) illustration of the binding pocket. This figure provides a 3D representation of B175L Cavity 4, the binding pocket binds with the top ligand, generated using the CD-Dock2 server. (B) 3D illustration of the ligand Benzaldehyde, 5-bromo-2-hydroxy-, (5-trifluoromethyl-2-pyridyl)hydrazone-B175L complex at Docking Pose. This figure provides a 3D representation of the Benzaldehyde, 5-bromo-2-hydroxy-, (5-trifluoromethyl-2-pyridyl)hydrazone ligand and B175L complex at the docking pose, generated using UCSF Chimera. (C) Two-dimensional (2D) interaction diagram of top ligand-protein complex, the interaction in green color represents conventional hydrogen bonding, while the blue line depicts halogen interaction between ligand and protein residues, the interaction in dark pink color represents Pi-Pi T-shaped bonds, and light pink lines depict alkali and Pi-alkali interactions.

**Table 3.** Noncovalent interactions of the selected ligand with protein B175L.

| Name | Distance | Category | Types |
| --- | --- | --- | --- |
| THR48 | 2.56965 | Hydrogen Bond | Conventional Hydrogen Bond |
| VAL46 | 2.45481 | Hydrogen Bond | Conventional Hydrogen Bond |
| PRO72 | 3.5427 | Halogen | Halogen (Fluorine) |
| PRO72 | 2.82304 | Halogen | Halogen (Fluorine) |
| PHE47 | 5.32418 | Hydrophobic | Pi-Pi T-shaped |
| VAL46 | 4.74154 | Hydrophobic | Alkyl |
| LEU43 | 4.13605 | Hydrophobic | Alkyl |
| PRO72 | 4.58338 | Hydrophobic | Alkyl |
| PHE74 | 4.86552 | Hydrophobic | Pi-Alkyl |
| PRO44 | 4.9547 | Hydrophobic | Pi-Alkyl |
| LYS52 | 5.35585 | Hydrophobic | Pi-Alkyl |
| PRO44 | 4.30411 | Hydrophobic | Pi-Alkyl |
| VAL54 | 5.24165 | Hydrophobic | Pi-Alkyl |
| VAL91 | 5.24664 | Hydrophobic | Pi-Alkyl |

The highest-affinity ligand localized to a cavity containing residues LEU43, PRO44, VAL46, PHE47, THR48, THR49, LYS52, TRP53, VAL54, PRO72, LYS73, PHE74, VAL91, GLY93, PHE128. Among them, 5 of the residues PRO72, LYS73, PHE74, VAL91, and GLY93 are within the zinc finger motif (60-104) of B175L (S1C Fig).

### MD simulations

To investigate the real-time dynamics and conformational stability of a protein upon binding to a specific ligand, 100.102 ns of MD simulations were performed on the best-docked compound (Benzaldehyde, 5-bromo-2-hydroxy-, (5-trifluoromethyl-2-pyridyl)hydrazone) at the protein binding site. Our analysis of simulation interaction diagrams (SIDs) for the 100.102 ns SPC water model-based simulations provided a better understanding of Protein RMSD, Ligand RMSD, Protein RMSF, Ligand RMSF, Protein-Ligand contacts, and ligand characteristics were analyzed.

#### RMSD of top ligand at B175L binding site

The RMSD of the B175L protein-ligand complex was determined over a 100.102 ns molecular dynamics simulation. Analysis of the protein backbone (Cα) and ligand RMSD was performed to evaluate the system’s stability. During the initial 20 ns of the simulation, the protein RMSD rapidly increased from approximately 2 Å to 14 Å. Subsequently, the protein RMSD fluctuated around an average of 13-14 Å for the remainder of the simulation period. Similarly, the ligand RMSD (fitted to the protein) also experienced a rapid increase in the first 20 ns, rising from approximately 1 Å to 12 Å. Following this initial phase, the ligand RMSD fluctuated and stabilized, averaging around 10-12 Å for the remaining 80 ns. The observed initial increase and subsequent stabilization in both protein and ligand RMSD suggest that while the complex undergoes conformational adjustments early in the simulation, it reaches a relatively stable state for the majority of the 100.102 ns trajectory (Fig 6).

**Fig 6.**
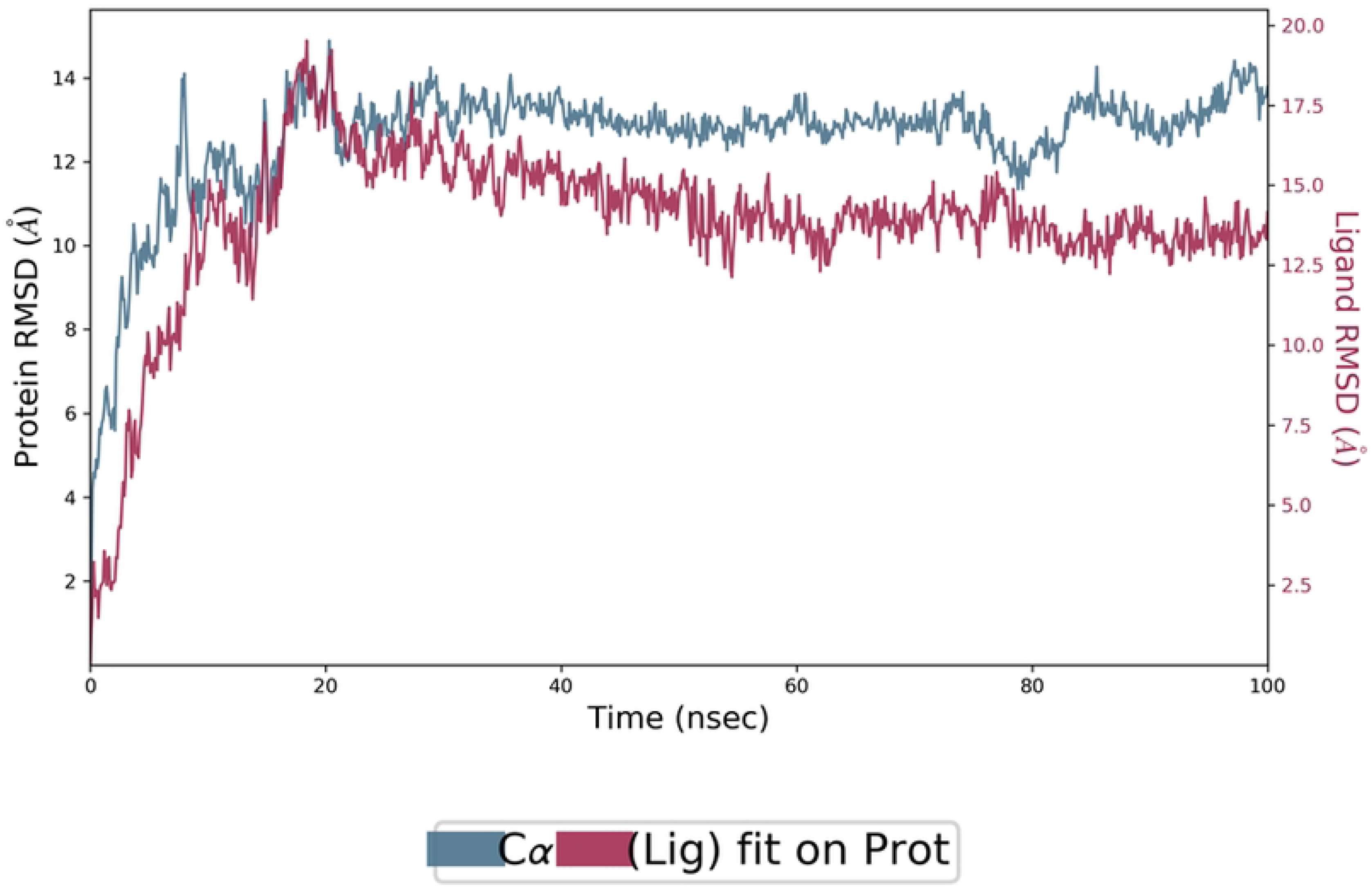
Protein-ligand root mean square deviation (RMSD) RMSD plot of B175L in complex with Benzaldehyde, 5-bromo-2-hydroxy-, (5-trifluoromethyl-2-pyridyl)hydrazone during the period of Molecular Dynamics simulations (MD Simulations). The blue line in the graph shows protein RMSD in the form of a C alpha chain of protein, and pink lines represent Lig fit Prot RMSD. Protein RMSD values are on the left y-axis, while lig fit prot RMSD values are on the right Y-axis. The X-axis shows the period of MD simulations in nanoseconds (nsec).

#### Protein and ligand RMSF

As shown in Fig 7A, the protein’s RMSF values ranged from approximately 0.5 Å to 8.5 Å, indicating varying degrees of flexibility across different residues. Residues with higher RMSF values are primarily located in loop regions, while those with lower values are found in more rigid secondary structure elements. The ligand RMSF values ranged from approximately 2.5 Å to 6.0 Å (Fig 7B). Fluctuations observed at ligand atoms 1 to 21 suggest dynamics within the binding pocket; however, no significant deviations were observed that would compromise the ligand’s overall stability or binding pose.

**Fig. 7.**
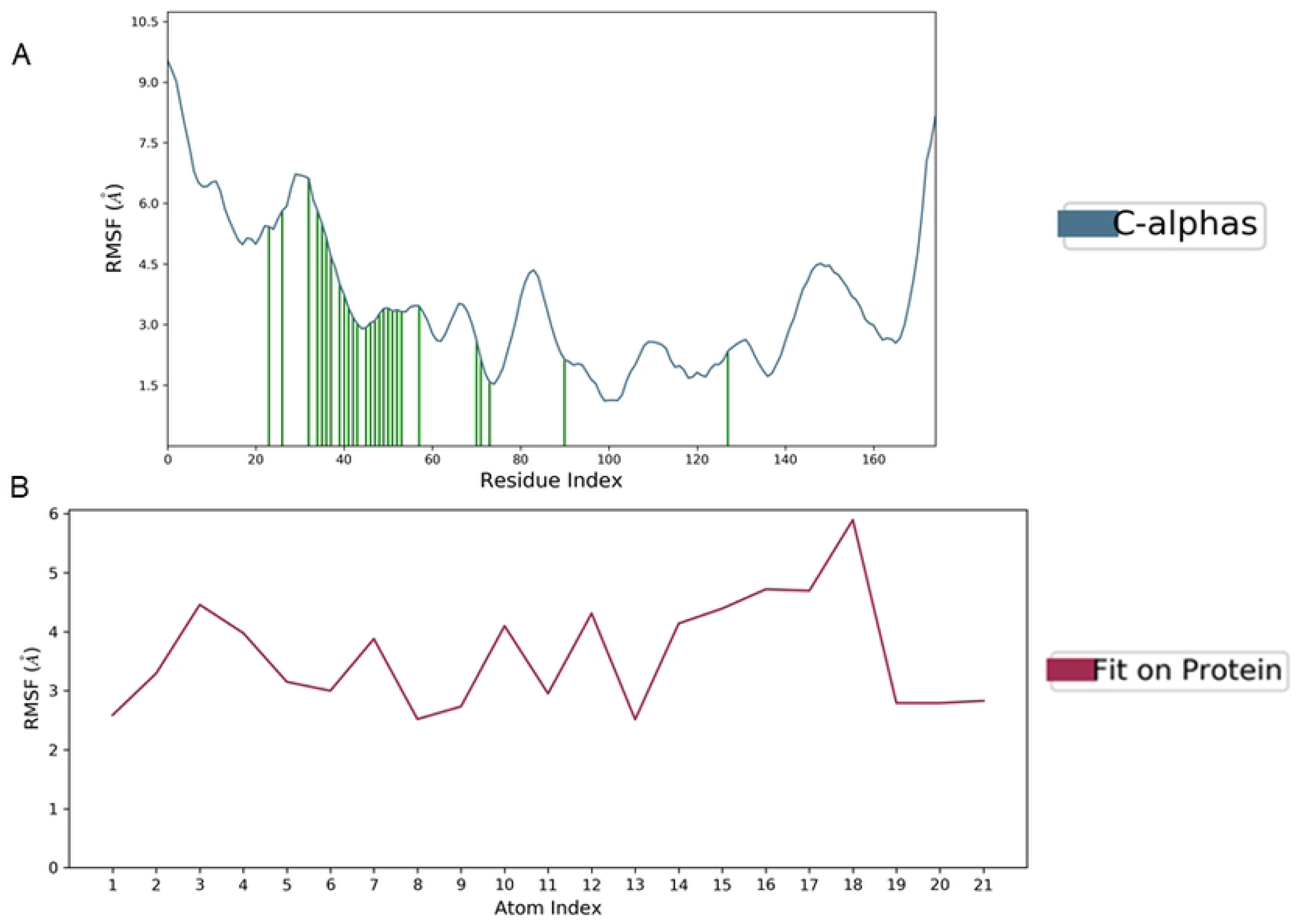
Protein and ligand root mean square fluctuations (RMSF) (A) RMSF plot of B175L where the active site residues are on the X-axis, RMSF values on the Y-axis, and the green bar indicates the point of contact of respective amino acid residues at the binding site with Benzaldehyde, 5-bromo-2-hydroxy-, (5-trifluoromethyl-2-pyridyl)hydrazone ligand. (B) Ligand RMSF. The pink line in the plot shows the RMSF of the ligand, the X-axis indicates residues of the ligand, and the Y-axis shows the value of RMSF.

#### Protein-ligand contacts

The ligand was in contact with 28 amino acid residues within the nucleotide-binding domain throughout the simulations. These residues formed critical interactions such as hydrogen bonds, hydrophobic contacts, and water bridges. Hydrogen bonds were observed consistently during the simulation, contributing significantly to the ligand’s stable binding. Several hydrophobic interactions were maintained, involving residues in the binding pocket that stabilize the ligand’s position. Water-mediated hydrogen bonds were detected between specific residues and the ligand, further enhancing interaction stability (Fig 8). The ligand interacted with PRO72, PHE74 residues in the zinc finger binding area of B175L with strong hydrophobic interactions.

**Fig 8.**
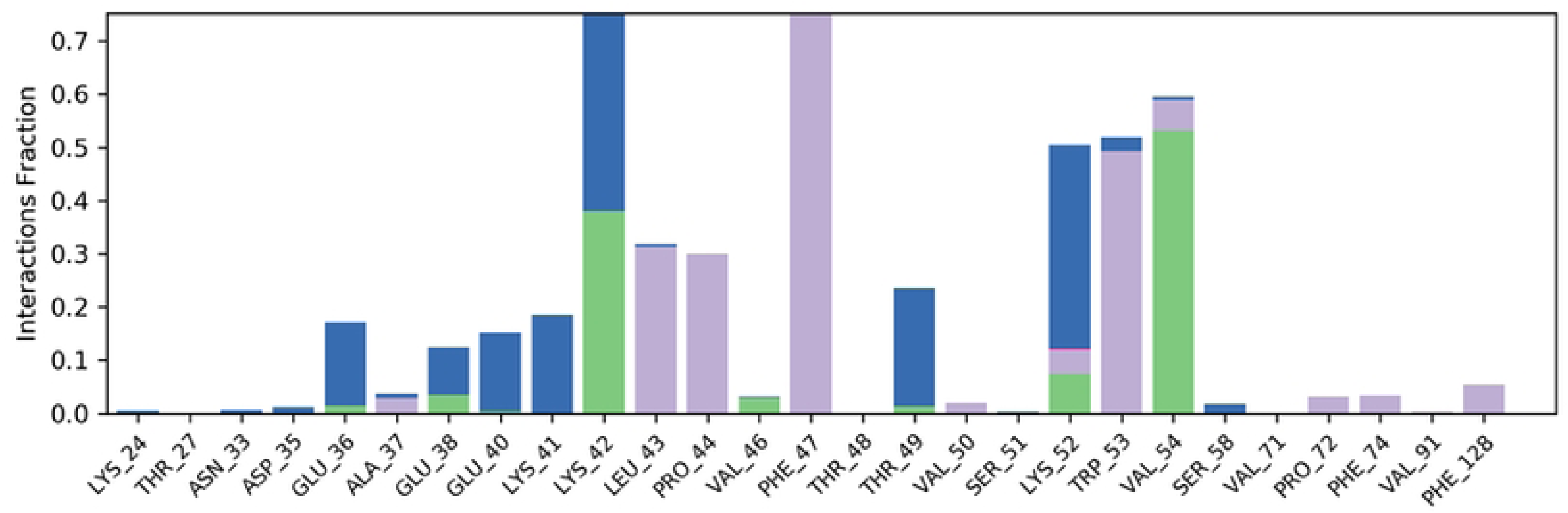
Protein-ligand contacts. This histogram shows ligand interaction with amino acids at the binding site, purple for hydrophobic interaction, green for hydrogen bond, pink for ionic interaction, and blue for water bridge. The X-axis indicates amino acid residues, while the Y-axis shows the Interaction fraction.

### Changes in the ligand (Benzaldehyde, 5-bromo-2-hydroxy-, (5-trifluoromethyl-2-pyridyl)hydrazone) properties

To assess the stability of the lead molecule in the binding sites of the B175L, the five molecular characteristics of the ligand (ligand RMSD, rGyr, MolSA, SASA, and PSAwere also investigated (S2 Fig).

#### Ligand RMSD

The ligand exhibited RMSD values that typically ranged from 0.2 Å to 1.6 Å, indicating a stable binding mode within the pocket (S2A Fig).

#### Radius of gyration

The rGyr of the ligand was found to be relatively constant, generally fluctuating between 4.2 Å and 4.8 Å, suggesting consistent compactness and shape throughout the simulation (S2B Fig).

#### Molecular surface area

The ligand’s MolSA was observed to be 260 Å² to 275 Å², indicating its overall size and surface characteristics (S2C Fig).

#### Solvent accessible surface area

The SASA of the ligand varied from approximately 40 Å² to 150 Å², demonstrating the extent to which the ligand was exposed to the solvent environment during the simulation (S2D Fig).

#### Polar surface area

The PSA of the ligand was found to range from 70 Å² to 95 Å², reflecting the contribution of polar atoms (oxygen and nitrogen) to the ligand’s overall surface area and its potential for polar interactions within the binding site (S2E Fig).

## Discussion

In this study, we described a new plant compound from *T. cordifolia* that aids in the inhibition of ASFV B175L by *in silico* pharmacological analysis. Our findings expand upon previous research on ASFV B175L immune evasion strategies (Ranathunga et al., 2023) and provide new insights into targets for vaccine development. The GC-MS analysis of *T. cordifolia* stem methanol extract revealed a diverse array of phytochemical compounds, with 124 identified. This aligns with previous studies highlighting the pharmacological potential of *T. cordifolia* (44). Our rigorous ADME/T screening process, adhering to Lipinski’s Rule of Five, identified 52 compounds with favorable drug-like properties and no predicted toxicity flags. This approach mirrors successful strategies used in other natural product-based drug discovery efforts. Our structural analysis and modeling of the ASFV B175L protein provided several important insights. The protein’s amino acid composition, with a high proportion of glutamic acid, serine, and threonine, suggests potential for post-translational modifications and regulatory functions (45). The predicted instability index and aliphatic index values provide clues about the protein’s stability and potential interaction in the cellular environment.

The AlphaFold3-generated model of B175L, refined using the GalaxyWeb server, demonstrated high confidence metrics and good stereochemical properties. The AlphaFold3 predicted structure was strictly validated using the confidence metrics outlined by the AlphaFold3 framework. According to the server’s guidelines, pTM evaluates the global structural accuracy of the entire complex, with values >0.5 indicating a plausible overall fold, while ipTM assesses the accuracy of subunit interfaces in multi-chain complexes, where scores >0.8 denote high confidence predictions and 0.6-0.8 in average confidence prediction (46) and raking score, which is internal confidence metrics >0.80 indicating high-quality prediction. This model allowed us to identify sites and structural characteristics that might be indicative of B175L function. A striking feature of B175L is a predicted disordered region, which constitutes 26% of the protein; intrinsically disordered regions play major roles in protein-protein interactions and regulation. The refined model was selected based on the optimization of several parameters. As detailed by (47), GalaxyWeb aims to balance various factors to achieve an optimal structure. A lower MolProbity score indicates better stereochemical quality, reflecting fewer steric clashes and improved geometry. A higher GDT-HA score signifies greater structural similarity to the native state, while a lower RMSD suggests a reduced deviation from the initial model. A lower clash score reflects fewer atomic clashes, and a lower number of poor rotamers improves side-chain conformations.

The structural validation of the refined B175L model was conducted using ProSA-web, PROCHECK, and MolProbity. The ProSA-web Z-score of -5.26 indicates that the overall fold is consistent with known protein structures (48). Ramachandran plot analysis revealed no residues in the disallowed region, confirming excellent stereochemical quality. The VERIFY3D score and ERRAT quality factor further supported the model’s reliability.

From our molecular docking studies, we identified Benzaldehyde, 5-bromo-2-hydroxy-, (5-trifluoromethyl-2-pyridyl)hydrazone as a promising lead compound, with high binding affinity to B175L. The formation of conventional hydrogen bonds and multiple hydrophobic interactions suggests a stable and specific binding mode. The MD simulations provided valuable insights into the stability and dynamics of the B175L-Benzaldehyde, 5-bromo-2-hydroxy-, (5-trifluoromethyl-2-pyridyl)hydrazone complex, revealing a stable interaction after initial conformational adjustments. The protein and ligand RMSD indicate a stable complex where initial fluctuations often reflect system equilibration before reaching stability. RMSF analysis showed varying flexibility across protein residues with higher values in loop regions, while the ligand’s RMSF suggested localized flexibility without compromising stability. The ligand formed consistent hydrogen bonds and hydrophobic interactions with 28 amino acid residues, underscoring its strong binding affinity. Notably, the presence of interacting residues is located within the zinc finger motif of B175L during MD simulations, which may be critical for its function. Additionally, ligand properties such as rGyr remained constant between 4.2 and 4.8, reflecting consistent compactness of Benzaldehyde, 5-bromo-2-hydroxy-, (5-trifluoromethyl-2-pyridyl)hydrazone within the binding pocket. MolSA and values indicated favorable surface characteristics for polar interactions and solvent accessibility, while SASA values demonstrated that Benzaldehyde, 5-bromo-2-hydroxy-, (5-trifluoromethyl-2-pyridyl)hydrazone was appropriately exposed to its environment while maintaining strong binding interactions.

These findings have important implications for our understanding of ASFV pathogenesis and the development of potential therapeutic strategies. By inhibiting B175L, our identification of small molecules effectively prevents B175L-mediated suppression of the IFN signaling pathway, restoring the host’s antiviral immune response. This restoration of the host’s antiviral immune response demonstrates B175L’s critical role in immune evasion and establishes a foundation for the development of effective ASFV drugs.

Future studies should focus on *in vitro* and *in vivo* studies, which are necessary to validate the efficacy of Benzaldehyde, 5-bromo-2-hydroxy-, (5-trifluoromethyl-2-pyridyl)hydrazone as an inhibitor of B175L function and its potential as an antiviral agent against AFSV.

## Conclusion

Based on the findings of this study, the *T. cordifolia-derived* Benzaldehyde, 5-bromo-2-hydroxy-, (5-trifluoromethyl-2-pyridyl)hydrazone possesses strong anti-ASFV B175L activity. Molecular Dynamics simulations studies were performed for 100.102 ns on a selected ligand, which provides clues regarding the structural stability of the protein-ligand complex. Our study combines computational approaches with natural product screening to identify a novel target and potential inhibitor of ASFV. This work not only advances our understanding of ASFV biology but also demonstrates the power of integrating traditional medicinal plant knowledge with modern computational drug discovery techniques.

## Supporting information

**S1 Fig. Graphical illustration of gas chromatography-mass spectrometry (GC-MS) chromatogram, modelled B175L, and zinc finger binding area of B175L**

(A) GC-MS chromatogram of *Tinospora cordifolia* stem methanol extraction. (B) Modeled B175L structure using the Alphafold3 server. The model is colored according to the predicted local distance difference test (pLDDT) score, a measure of the model’s confidence in the predicted structure. Regions colored blue indicate very high confidence (pLDDT > 90), cyan indicates confident regions (90 > pLDDT >70), yellow indicates low confidence (70 > pLDDT > 50), and orange indicates very low confidence (pLDDT < 50). (C) Zinc finger binding area of B175L, referring to NCBI Conserved Domain. The binding region of B175L is located between amino acid residues 60 and 104 within its 175-amino-acid sequence. Amino acids located in the docking cavity of the L1 ligand are shown in red.

**S2 Fig. Changes in the ligand properties during simulation**

(A) Ligand root mean square deviation (RMSD) plot. The Y-axis indicates the RMSD value of the ligand in Angstrom (Å). (B) Radius of gyration (rGyr) graph. The Y-axis shows rGyr in Å. (C) Molecular Surface Area (MolSA) graph. The Y-axis indicates MolSA values of ligands in 2 Å. (D) Solvent-accessible surface area (SASA) plot. The Y-axis shows SASA values of the ligand in 2 Å. (E) Polar surface area (PSA) graph. The Y-axis indicates the PSA values of the ligand in 2 Å. The X-axis of all these 4 graphs indicates the time in nanoseconds (nsec).

## Notes

### Competing Interest Statement

The authors have declared no competing interest.

## References

1. Urbano AC, Ferreira F. African swine fever control and prevention: an update on vaccine development. Emerging Microbes and Infections [Internet]. 2022;11(1):2021–33. Available from: 10.1080/22221751.2022.2108342

2. Juszkiewicz M, Walczak M, Woźniakowski G, Szczotka-Bochniarz A. Virucidal activity of plant extracts against african swine fever virus. Pathogens [Internet]. 2021;10(11). Available from: 10.3390/pathogens10111357

3. Organisation W, Health A, January S. AFRICAN SWINE FEVER (ASF) Situation Report 60 Period. 2024;(May):1–7.

4. Ranathunga L, Abesinghe S, Cha JW, Dodantenna N, Chathuranga K. Inhibition of STING - mediated type I IFN signaling by African swine fever virus DP71L. Veterinary Research [Internet]. 2025;1–16. Available from: 10.1186/s13567-025-01474-3

5. Kim YJ, Park B, Kang HE. Control measures to African swine fever outbreak: active response in South Korea, preparation for the future, and cooperation. Journal of Veterinary Science [Internet]. 2021;22(1):1–14. Available from: 10.4142/jvs.2021.22.e13

6. Cruz C, Pineiro M, Moreno R. Eur 31718 En [Internet]. 2023. Available from: 10.2760/970562

7. Kaushik N, Patel P, Gupta R, Jaiswal A, Negi M, Borkar SB, et al. Eco-friendly materials for next-generation vaccination: From concept to clinical reality. SmartMat [Internet]. 2024; Available from: 10.1002/smm2.1274

8. AL-Ballawi ZFS, Redhwan NA, Ali M. In Vitro Studies of Some Medicinal Plants Extracts for Antiviral Activity against Rotavirus. IOSR Journal of Pharmacy and Biological Sciences [Internet]. 2017;12(02):53–8. Available from: 10.9790/3008-1202025358

9. Brogi S, Ramalho TC, Kuca K, Medina-Franco JL, Valko M. Editorial: In silico Methods for Drug Design and Discovery. Frontiers in Chemistry [Internet]. 2020;8(August):1–5. Available from: 10.3389/fchem.2020.00612

10. Scope T, Tinospora OF, In C. THERAPEUTIC SCOPE OF TINOSPORA CORDIFOLIA IN MODERN. 2025;14(3):382–407. Available from: 10.20959/wjpr20253-35419

11. Ranathunga L, Dodantenna N, Cha J-W, Chathuranga K, Chathuranga WAG, Subasinghe A, et al. pathway by targeting STING and 2 ′ 3 ′ -cGAMP. 2023;97(October).

12. Aly SH, Uba AI, Nilofar N, Majrashi TA, El Hassab MA, Eldehna WM, et al. Chemical composition and biological activity of lemongrass volatile oil and n-Hexane extract: GC/MS analysis, in vitro and molecular modelling studies. PLoS ONE [Internet]. 2025;20(2 February):1–19. Available from: 10.1371/journal.pone.0319147

13. Tong K, Dai L, Rui W, Zhang Y, Fu J, Liao Y, et al. GC-MS, LC-MS, and network pharmacology analysis to investigate the chemical profiles and potential pharmacological activities in flower buds and flowers of Lonicera japonica Thunb. PLoS ONE [Internet]. 2025;20(4 April):1–19. Available from: 10.1371/journal.pone.0320293

14. Ghosh G, Panda P, Rath M, Pal A, Sharma T, Das D. GC-MS analysis of bioactive compounds in the methanol extract of clerodendrum viscosum leaves. Pharmacognosy Research [Internet]. 2015;7(1):110–3. Available from: 10.4103/0974-8490.147223

15. Srikalyani V, Ilango K. Chemical fingerprint by HPLC-DAD-ESI-MS, GC-MS analysis and anti-oxidant activity of Manasamitra Vatakam: A herbomineral formulation. Pharmacognosy Journal [Internet]. 2020;12(1):115–23. Available from: 10.5530/pj.2020.12.18

16. Sangma C, Chetia D, Borthakur M, Patowary L, Tayeng D. In-silico design and screening of cephalosporin derivatives for their inhibitory potential against Haemophilus influenza. Sciences of Phytochemistry [Internet]. 2022;1(2):1–10. Available from: 10.58920/sciphy01020001

17. Boudou F, Belakredar A, Keziz A, Alsaeedi H, Cornu D, Bechelany M, et al. Camellia sinensis phytochemical profiling, drug-likeness, and antibacterial activity against gram-positive and gram-negative bacteria: in vitro and in silico insights. Frontiers in Chemistry [Internet]. 2025;13(March):1–17. Available from: 10.3389/fchem.2025.1555574

18. Ibrahim ZY, Uzairu A, Shallangwa GA, Abechi SE. Application of QSAR Method in the Design of Enhanced Antimalarial Derivatives of Azetidine-2-carbonitriles, their Molecular Docking, Drug-likeness, and SwissADME Properties. Iranian Journal of Pharmaceutical Research [Internet]. 2021;20(3):254–70. Available from: 10.22037/ijpr.2021.114536.14901

19. Boudou F, Belakredar A, Berkane A, Keziz A, Alsaeedi H, Cornu D, et al. Phytochemical profiling and in silico evaluation of Artemisia absinthium compounds targeting Leishmania N-myristoyltransferase: molecular docking, drug-likeness, and toxicity analyses. Frontiers in Chemistry [Internet]. 2024;12(November):1–16. Available from: 10.3389/fchem.2024.1508603

20. Kadioglu O, Klauck SM, Fleischer E, Shan L, Efferth T. Selection of safe artemisinin derivatives using a machine learning-based cardiotoxicity platform and in vitro and in vivo validation. Archives of Toxicology [Internet]. 2021;95(7):2485–95. Available from: 10.1007/s00204-021-03058-4

21. Singh N, Sharma P, Pal MK, Kahera R, Badoni H, Pant K, et al. Computational drug discovery of phytochemical alkaloids targeting the NACHT/PYD domain in the NLRP3 inflammasome. Scientific Reports [Internet]. 2025;15(1):1–23. Available from: 10.1038/s41598-024-79054-2

22. Kaur D, Patiyal S, Arora C, Singh R, Lodhi G, Raghava GPS. In-Silico Tool for Predicting, Scanning, and Designing Defensins. Frontiers in Immunology [Internet]. 2021;12(November):1–12. Available from: 10.3389/fimmu.2021.780610

23. Sanjida S, Mou MJ, Islam SI, Sarower-E-Mahfuj M. An In-silico approaches for identification of potential natural antiviral drug candidates against Erythrocytic necrosis virus (Iridovirus) by targeting Major capsid protein: A Quantum mechanics calculations approach. International Journal of Life Sciences and Biotechnology [Internet]. 2022;5(3):294–315. Available from: 10.38001/ijlsb.1074392

24. Of A, Proteins S, Breast OF, By P, Bioinformatics U. ANALYSIS OF SEROLOGICAL PROTEINS OF BREAST CANCER. 2020;3(4):187–94. Available from: 10.31580/pjmls.v3i4.1730

25. Das SC, Biswas S, Khan O, Akter R, Azad AK, Sarkar SK, et al. Evaluation of anti-inflammatory and wound healing properties of Tinospora cordifolia extract. 2025;1–14. Available from: 10.1371/journal.pone.0317928

26. Abramson J, Adler J, Dunger J, Evans R, Green T, Pritzel A, et al. Accurate structure prediction of biomolecular interactions with AlphaFold 3. Nature [Internet]. 2024;630(8016):493–500. Available from: 10.1038/s41586-024-07487-w

27. Islam SI, Sanjida S, Mou MJ, Sarower-E-Mahfuj M, Nasir S. In-silico functional annotation of a hypothetical protein from Edwardsiella tarda revealed Proline metabolism and apoptosis in fish. International Journal of Life Sciences and Biotechnology [Internet]. 2022;5(1):78–96. Available from: 10.38001/ijlsb.1032171

28. Hosen MI, Mia ME, Islam MN, Khatun MUS, Emon TH, Hossain MA, et al. In-silico approach to characterize the structure and function of a hypothetical protein of Monkeypox virus exploring Chordopox-A20R domain-containing protein activity. Antiviral Therapy [Internet]. 2024;29(3). Available from: 10.1177/13596535241255199

29. Oladejo D, Oduselu G, Dokunmu T, Isewon I, Okafor E, Iweala EEJ, et al. In silico evaluation of inhibitors of Plasmodium falciparum AP2-I transcription factor. The FASEB Journal [Internet]. 2022;36(S1):1–16. Available from: 10.1096/fasebj.2022.36.s1.l7455

30. Singh R, Pokle AV, Ghosh P, Ganeshpurkar A, Swetha R, Singh SK, et al. Pharmacophore-based virtual screening, molecular docking and molecular dynamics simulations study for the identification of LIM kinase-1 inhibitors. Journal of Biomolecular Structure and Dynamics [Internet]. 2023;41(13):6089–103. Available from: 10.1080/07391102.2022.2101529

31. Pundir H, Pathak R, Pant T, Pant M, Chandra S, Tamta S. In Silico Screening of Phytochemicals Targeting SmdCD of Streptococcus mutans using Molecular Docking Approach. Trends in Sciences [Internet]. 2023;20(6). Available from: 10.48048/tis.2023.6036

32. Luthufi H, Mudalige H, Perera O. Protein-Ligand Docking To Identify Potential Phytochemicals Against Breast Cancer Protein Receptor Using Autodock. Gari International Journal of Multidisciplinary Research. 2022;8(4):108–27.

33. Kumar P, Alpana A, Dinesh C, Abhinav B. GSC Biological and Pharmaceutical Sciences In silico screening and molecular docking of bioactive agents towards human coronavirus receptor. 2020;11(01):132–40.

34. Amraei S, Ahmadi S. Nano Micro Biosystems. 2022;1(1):22–6.

35. Azmal M, Paul JK, Prima FS, Talukder OF, Ghosh A. An in silico molecular docking and simulation study to identify potential anticancer phytochemicals targeting the RAS signaling pathway. PLoS ONE [Internet]. 2024;19(9 September):1–27. Available from: 10.1371/journal.pone.0310637

36. Elzupir AO. Molecular docking and dynamics investigations for identifying potential inhibitors of the 3-chymotrypsin-like protease of sars-cov-2: Repurposing of approved pyrimidonic pharmaceuticals for covid-19 treatment. Molecules [Internet]. 2021;26(24). Available from: 10.3390/molecules26247458

37. Iqbal D, Alsaweed M, Jamal QMS, Asad MR, Rizvi SMD, Rizvi MR, et al. Pharmacophore-Based Screening, Molecular Docking, and Dynamic Simulation of Fungal Metabolites as Inhibitors of Multi-Targets in Neurodegenerative Disorders. Biomolecules [Internet]. 2023;13(11). Available from: 10.3390/biom13111613

38. Akash S, Islam MR, Bhuiyan AA, Islam MN, Bayıl I, Saleem RM, et al. In silico evaluation of anti-colorectal cancer inhibitors by Resveratrol derivatives targeting Armadillo repeats domain of APC: molecular docking and molecular dynamics simulation. Frontiers in Oncology [Internet]. 2024;14(April):1–15. Available from: 10.3389/fonc.2024.1360745

39. Dobson L, Reményi I, Tusnády GE. CCTOP: A Consensus Constrained TOPology prediction web server. Nucleic Acids Research [Internet]. 2015;43(W1):W408–12. Available from: 10.1093/nar/gkv451

40. Manandhar S, Sankhe R, Priya K, Hari G, Kumar B H, Mehta CH, et al. Molecular dynamics and structure-based virtual screening and identification of natural compounds as Wnt signaling modulators: possible therapeutics for Alzheimer’s disease. Molecular Diversity [Internet]. 2022;26(5):2793–811. Available from: 10.1007/s11030-022-10395-8

41. Vemula D, Maddi DR, Bhandari V. Homology modeling, virtual screening, molecular docking, and dynamics studies for discovering Staphylococcus epidermidis FtsZ inhibitors. Frontiers in Molecular Biosciences [Internet]. 2023;10(March):1–24. Available from: 10.3389/fmolb.2023.1087676

42. Parthiban S, Raja P, Parthiban M, Yamini C. Tackling African swine fever in India. CAB Rev Perspect Agric Vet Sci Nutr Nat Resour [Internet]. 2023 Dec 22; Available from: 10.1079/cabireviews.2023.0050

43. Dr.Kamble P. S. SPVDACK. Ijfans International Journal of Food and Nutritional Sciences. I) Journal. 2022;11(10):3870–7.

44. Chester K, Zahiruddin S, Ahmad A, Khan W, Paliwal S, Ahmad S. Bioautography-based Identification of Antioxidant Metabolites of Solanum nigrum L. and Exploration Its Hepatoprotective Potential agChester, K. et al. (2017) ‘Bioautography-based Identification of Antioxidant Metabolites of Solanum nigrum L. and Explorati. Pharmacognosy Magazine [Internet]. 2017;13 (Suppl(62):179–88. Available from: 10.4103/pm.pm

45. Click M. Manuscript Click here to view linked References. Brain [Internet]. 2010;2(i):617–38. Available from: 10.1016/j.actpsy.2011.12.005

46. Xu J, Zhang Y. How significant is a protein structure similarity with TM-score = 0.5? Bioinformatics [Internet]. 2010;26(7):889–95. Available from: 10.1093/bioinformatics/btq066

47. Ko J, Park H, Heo L, Seok C. GalaxyWEB server for protein structure prediction and refinement. Nucleic Acids Research [Internet]. 2012;40(W1):294–7. Available from: 10.1093/nar/gks493

48. Wiederstein M, Sippl MJ. ProSA-web: Interactive web service for the recognition of errors in three-dimensional structures of proteins. Nucleic Acids Research [Internet]. 2007;35(SUPPL.2):407–10. Available from: 10.1093/nar/gkm290

